# Unifying community-wide whole-brain imaging datasets enables robust automated neuron identification and reveals determinants of neuron positioning in *C. elegans*

**DOI:** 10.1101/2024.04.28.591397

**Authors:** Daniel Y. Sprague, Kevin Rusch, Raymond L. Dunn, Jackson M. Borchardt, Steven Ban, Greg Bubnis, Grace C. Chiu, Chentao Wen, Ryoga Suzuki, Shivesh Chaudhary, Hyun Jee Lee, Zikai Yu, Benjamin Dichter, Ryan Ly, Shuichi Onami, Hang Lu, Koutarou D. Kimura, Eviatar Yemini, Saul Kato

## Abstract

We develop a data harmonization approach for *C. elegans* volumetric microscopy data, still or video, consisting of a standardized format, data pre-processing techniques, and a set of human-in-the-loop machine learning based analysis software tools. We unify a diverse collection of 118 whole-brain neural activity imaging datasets from 5 labs, storing these and accompanying tools in an online repository called WormID (wormid.org). We use this repository to train three existing automated cell identification algorithms to, for the first time, enable accuracy in neural identification that generalizes across labs, approaching human performance in some cases. We mine this repository to identify factors that influence the developmental positioning of neurons. To facilitate communal use of this repository, we created open-source software, code, web-based tools, and tutorials to explore and curate datasets for contribution to the scientific community. This repository provides a growing resource for experimentalists, theorists, and toolmakers to (a) study neuroanatomical organization and neural activity across diverse experimental paradigms, (b) develop and benchmark algorithms for automated neuron detection, segmentation, cell identification, tracking, and activity extraction, and (c) inform models of neurobiological development and function.

## Introduction

Whole-brain imaging experiments with single-neuron resolution (herein shortened to simply “whole-brain imaging”) have undergone explosive growth since first demonstrated in the nematode *C. elegans,* a millimeter-sized worm, and the zebrafish *D. rerio* in 2013.^1,2^ Since then, these methods have been widely adopted and advanced in the worm^3,4,5,6,7^, zebrafish^8,9,10,11,12,13^, and larval^14^ and adult^15,16^ fly communities. Moreover, there have been significant efforts and advances in neuron-resolution imaging of multiple and/or large brain regions in mammals, rapidly approaching whole-brain imaging, especially in mice.^17,18,19,20,21^

In *C. elegans*, whole-brain imaging datasets have enabled characterization of neural dynamics^3,6^, functional connectivity^26,22,23,24^, and the roles of individual neurons during behavior^7^. These studies leverage the property of eutely in this organism: each cell has a unique and stereotyped identity, consistent across every animal, that allows for data from individual neurons to be pooled and compared across multiple trials and animals. However, analyses of these experiments are bottlenecked by the need to determine the unique identities of each neuron in 3D volumetric recordings. Manual cell identification from fluorescent microscopy imagery is a notoriously difficult skill, requiring substantial expertise and labor. This task is particularly difficult for neurons labeled with nuclear localized fluorophores, which is typical for whole-brain recordings. We recently developed NeuroPAL^6^, the first method where the unique identity of every single neuron can be distinguished by an invariant fluorescent color barcode in living animals at all developmental stages of both sexes.^25^ NeuroPAL has greatly simplified the task of cell identification and has thus seen rapid adoption, with at least 6 labs^6,7,26,22,24^ publishing whole-brain imaging datasets using these animals, and many more labs incorporating the system into their experimental protocols since its release in 2021.

Despite this innovation, neural identification remains a challenging task that requires expertise and many hours of manual work. In the past few years, researchers have proposed various algorithmic auto-identification approaches to attack this problem.^28,29,26,30,31,32^ However, none of them have achieved widespread adoption, due at least in part to their incompatibility with different microscopy data formats and low performance on data acquired from different labs. Automatic approaches to the complementary problem of tracking neurons across video frames have achieved some generalized performance across various datasets^33,34^, but so far there have not been efforts to perform similar training and benchmarking for automatic cell identification. In order to build automatic approaches that are robust, accurate, and generalizable, there is a critical need for a standardized format and compatible tools trained and benchmarked on a consolidated corpus of data that reflects the heterogeneity of microscopy equipment, experimental conditions, and protocols across labs.

To address this need, we take a data harmonization approach: a process of combining datasets from different sources and homogenizing them to produce a substantially larger data corpus that, in our case, minimizes non-biological inconsistencies across individual datasets while increasing the overall biological diversity of training and benchmarking data. Harmonization includes: i) aggregating the data, ii) converting it to a standardized format, iii) normalizing it, iv) handling duplicate and missing data, and v) pre-processing data to register it to a common space and coordinate system. Data harmonization is standard in many data science fields but has seen slower adoption in the life sciences^35^. Similar efforts to standardize data formats and build large corpuses of data have been essential in the development and benchmarking of many modern machine learning algorithms.^36,37,38^

We introduce WormID (wormid.org). This resource consists of: i) data harmonization tools including a standardized file format for both raw and processed data alongside related metadata that extends the existing Neurodata Without Borders (NWB) format, ii) pre-processing to align the color and coordinate space of new datasets, and iii) open-source software to analyze whole-brain activity images. We also provide tutorials and documentation that enable researchers to easily incorporate these tools into their data pipelines. Finally, we provide a large online corpus of harmonized *C. elegans* whole-brain activity imaging and structural data that can be used for large-scale experimental analysis, neurobiological modeling, and algorithmic development. This corpus is stored in a popular community archive called the Distributed Archives for Neurophysiology Data Integration (DANDI), which serves as a repository for experimental neuroscience data from a variety of model organisms.^39^

By aggregating a diversity of datasets from multiple labs into a large data corpus, we achieve a substantial boost in the performance of three existing neural auto-identification algorithms, arguably moving into the regime of practical utility for the broader community of users. Furthermore, we mine this corpus to investigate the relationship between neural lineage, synaptic connectivity, and somatic positioning of *C. elegans* neurons to better understand the factors that drive the positioning of neurons in the adult worm.

This corpus and set of tools should be of wide utility to *C. elegans* researchers. We hope it will serve as a seed for continued community aggregation of brain imaging datasets and further the development and improvement of community data-analysis tools applicable across many model organisms.

## Results

### A standardized format for whole-brain *C. elegans* recordings enables data aggregation and algorithm interoperability

Current state-of-the-art whole-brain recordings of *C. elegans* typically consist of a combination of structural images that often use the NeuroPAL multi-channel fluorescent system to determine neuron identities (**Fig. 1a-b**) and time series images of neural activity acquired by using genetically-encoded activity sensors (e.g., GCaMP6s^40^) (**1c**). This imaging is performed either on immobilized worms (often constrained within a microfluidic chip to maximize image quality^1,3,41^) or on freely-moving worms.^4,5^ To aid interpretation, herein we visualize whole-brain structural NeuroPAL images via i) an unrolled ‘butterfly’ plot of neuron positions that projects the 3D worm structure into a 2D plane (**Fig. 1a**), ii) a 2D projection plot of the NeuroPAL color space (**Fig. 1b**), and iii) 2D dorsal-ventral and lateral projection plots of the neurons (**Fig. 1b**). These visualizations facilitate quick comparisons of neuron color and position from different samples and fine-tuning of their global alignment.

**Figure 1:**
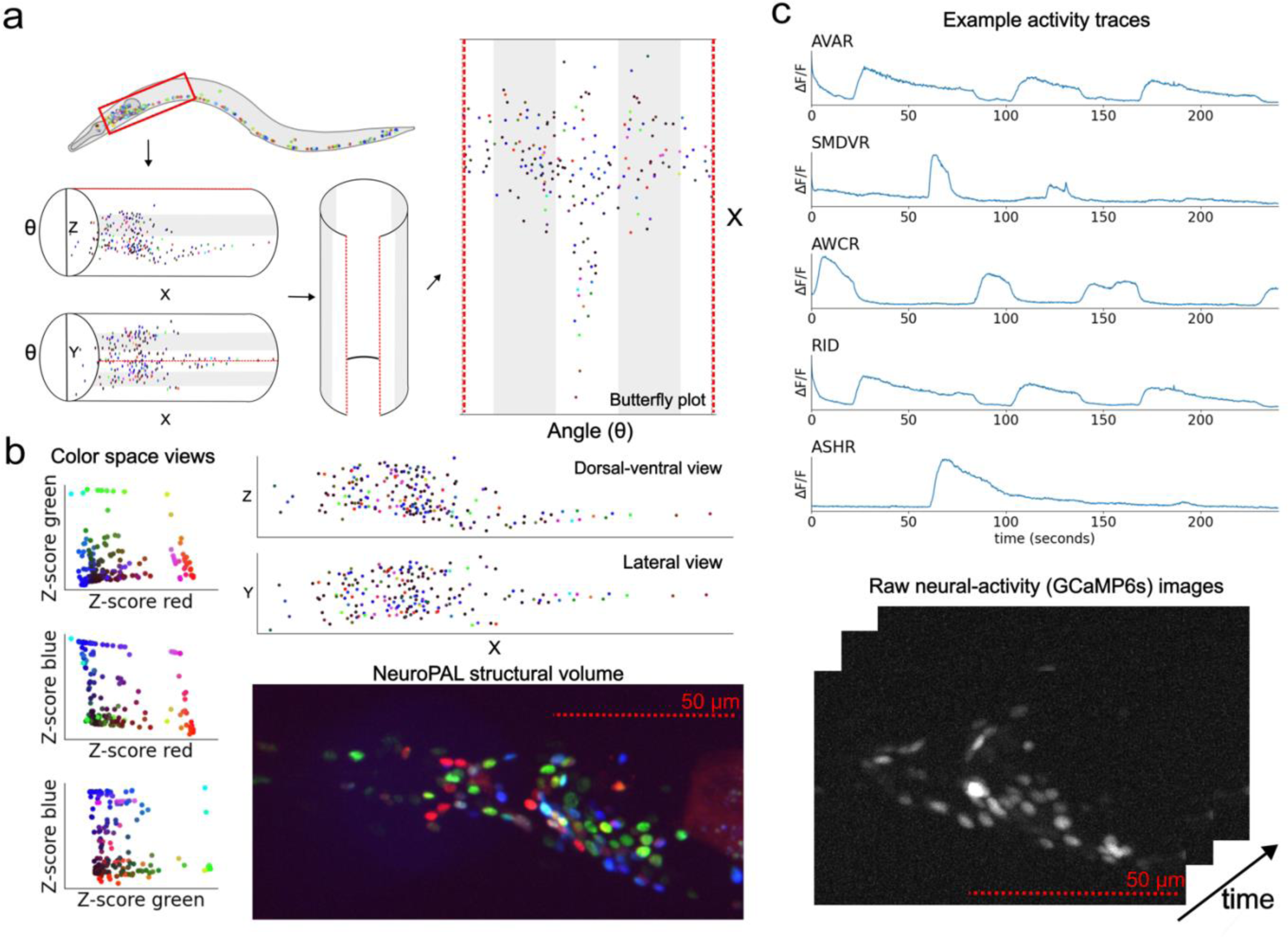
NWB file contents. (a) Illustration of a NeuroPAL worm. The head is highlighted by a red box. A butterfly plot visualizes the full 2D representation of the worm brain by projecting neurons onto the surface of the cylindrical body and then unrolling the cylinder. Neuron centers are colored using their composite NeuroPAL expression. (b) Visualizations of the raw NeuroPAL structural image and 2D projections of its RG, RB, GB color subspaces and XZ and XY projections of its neuron positions. Neuron centers are colored using their composite NeuroPAL expression. (c) Example activity traces for five neurons contained in the NWB file and of the raw neural-activity (GCaMP6s) images.

All associated raw data and metadata is stored in the standardized NWB^42^ file format with an additional extension that we developed, *ndx-multichannel-volume* (ndx = neurodata extension), to provide support for multi-channel volumetric recordings and *C. elegans* specific metadata (**Fig. 2a**). This extension is available in the NWB Extensions Catalog and is now the official NWB standard for data sharing of *C. elegans* whole-brain neural-activity imaging. NWB data is hierarchically organized with basic metadata stored at the file’s root level, raw data stored in the ‘acquisition module’, and various processed experimental data stored in ‘processing modules’ (**Fig. 2b**). Individual NWB files contain a single experimental run for a single animal. These NWB files are then stored and accessed from the DANDI archive, where they receive a unique persistent digital object identifier (DOI) in accordance with the International Organization for Standardization (ISO).

**Figure 2:**
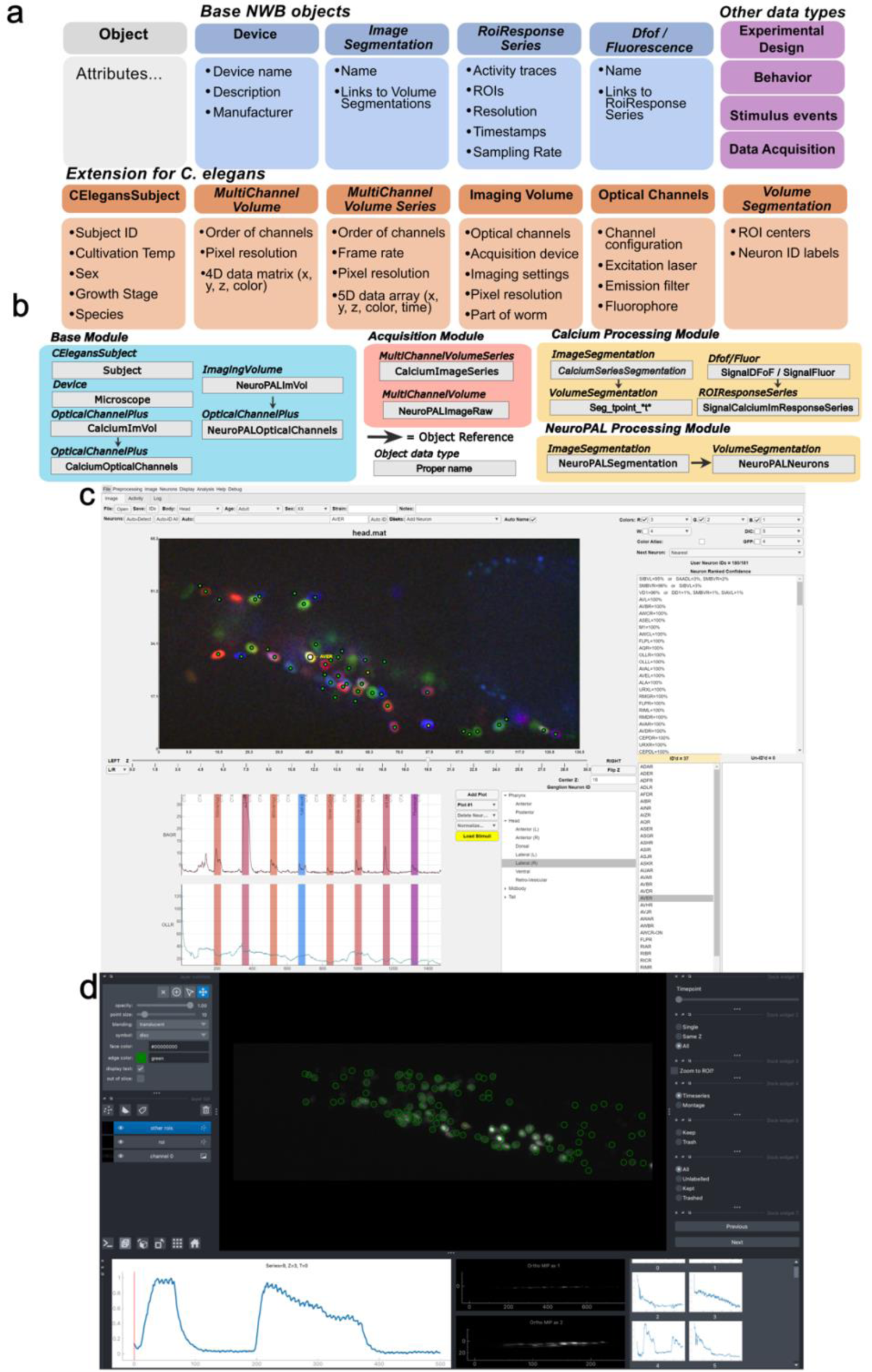
NWB schema and two software programs with NWB I/O support. (a) Names and content for objects used in *C. elegans* optophysiology NWB files. (b) File organization hierarchy of NWB files for *C. elegans* optophysiology. Modules are structured like folders within the root file in an HDF5-based hierarchy. (c-d) NeuroPAL ID software (c) and eats-worm software (d) GUIs with NWB I/O support for visualization and annotation of NeuroPAL structural images, neural segmentation and automated ID, and time-series of neural-activity with stimulus-presentation in immobilized worms.

We incorporated NWB *ndx-multichannel-volume* read and write functionality into two software tools. These independent software implementations both offer both user-friendly GUIs that analyze *C. elegans* NeuroPAL structural images and neural activity in immobilized worms (NeuroPAL software https://github.com/Yemini-Lab/NeuroPAL_ID, **Fig. 2c** and eats-worm software https://github.com/focolab/eats-worm, **Fig. 2d**). This functionality can be straightforwardly incorporated into other data-analysis pipelines and software.

In **Table 1** we present a summary of the data we aggregated and harmonized into a corpus: 108 worms from six datasets acquired by five different labs, each with segmented neurons and human-labeled identities. This corpus can be mined for biological insights, training and benchmarking of machine-vision approaches, and neurobiological studies of structural and neural-activity time-series data. Each of these datasets is stored on DANDI and range from a few hundred megabytes to several terabytes (see **Methods** for dataset references). DANDI supports streaming from the cloud and allows users to selectively load data objects and data chunks, substantially reducing the local data storage and RAM requirements necessary to work with this data on a personal computer.

**Table 1:** Summary of aggregated dataset characteristics. NP dataset comes from the original NeuroPAL paper^6^, obtained as part of a collaboration between the labs of Oliver Hobert (Columbia University) and Aravinthan D.T. Samuel (Harvard University).

| Dataset # | # of Worms in the Dataset | Lab Code | NeuroPAL, GCaMP, or Both | # of Segmented Neurons (Avg) | # of ID Labels (Avg) |
| --- | --- | --- | --- | --- | --- |
| NP | 10 | NP | NeuroPAL | 189-196 (193) | 186-193 (190) |
| 1 | 21 | EY | Both | 166-188 (177) | 164-184 (175) |
| 2 | 9 | HL | NeuroPAL | 113-125 (119) | 58-69 (64) |
| 3 | 9 | KK | Both | 149-163 (154) | 149-163 (154) |
| 4 | 38 | SF | Both | 29-96 (70) | 29-96 (70) |
| 5 | 21 | SK1 | Both | 78-139 (111) | 30-82 (48) |
| 6 | 10 | SK2 | NeuroPAL | 166-180 (173) | 38-63 (49) |
| Summary | 118 | 5 labs |  | 29-196 (126) | 29-193 (99) |

### An updated atlas of the *C. elegans* hermaphrodite head

Our multi-lab data corpus allows data scientists to train and benchmark the performance of algorithms for automated neuron identification using datasets that reflect real-world diversity. In this section, we focus on the statistical atlas approach presented in Varol et al. 2020.^28^ This approach was the first to take advantage of the color information provided by NeuroPAL and was presented alongside the original NeuroPAL work.^6^ This neuron identification assignment algorithm was framed as a bipartite graph matching problem, with the goal of minimizing the total assignment cost using the well-known Hungarian algorithm.^43^ Cost is calculated by comparing neuron position and color in the animal sample with the mean and covariance of neuron position and colors in a reference statistical atlas (see **Methods**). The original atlas presented in the paper was trained on 10 worms from the original NeuroPAL work. We retrain this atlas using the full multi-lab corpus that we present in this work, increasing the training set by over 10-fold. The statistical atlas generated by this approach serves the additional purpose of characterizing the mean and covariance of neuron position and color across the whole corpus of data.

In **Fig. 3** we present visualizations of the statistical atlas of neuron colors and positions trained on 104 of the 118 worms in our consolidated NWB/DANDI dataset as well as on the smaller dataset of 10 worms used in the original NeuroPAL paper (employing the Statistical Atlas algorithm in Yemini et al. 2021 and Varol et al. 2020^6,28^). 14 worms were omitted from the atlas due to large nonlinear deformities or obvious artifacts. With 104 worms, this represents, to the best of our knowledge, the most broadly trained statistical atlas for *C. elegans* neuron positions and NeuroPAL coloring available. By leveraging the diversity of the multi-lab corpus this atlas captures variability between individual worms, strains, and lab-specific experimental conditions. This atlas can be used as a basis for automatic labeling algorithms and biological investigations of neuron positions and brain organization. This statistical atlas further complements detailed electron-microscopy (EM) based anatomical atlases with cellular structural detail and provides nearly 100 more animals in its corpus than the approximately 10 EM ones available.^44,45,46^ Although our corpus lacks the synaptic connectivity found in the EM datasets, it provides the complementary functional activity that is not available from EM imaging.

**Figure 3.**
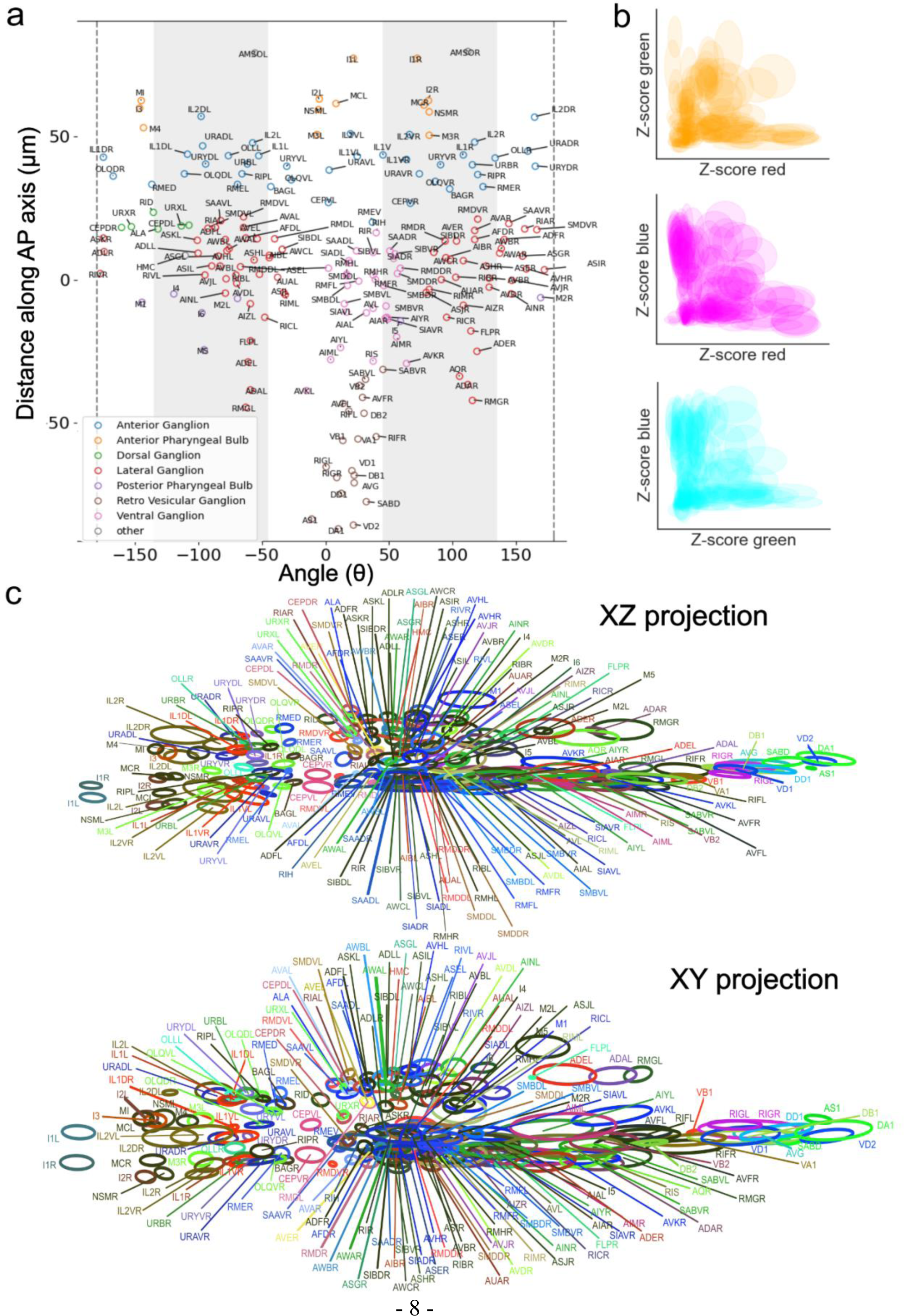
Multi-lab atlas of *C. elegans* neurons and their positional variability. (a) Butterfly plot showing the mean locations of neurons in the atlas colored by ganglion. (b) 2D color plots showing the distribution of neuron colors in the atlas. (b-c) Ellipses represent covariance (1 SD) and are centered at the mean for each neuron. (c) 2D projections of neuron positions and colors in the aligned atlas space. XZ projection (top) and XY projection (bottom). Ellipses are colored by the mean color, per color channel, for each neuron in the atlas.

WormID.org supplies links to the software, visualization tools, and datasets discussed previously in this paper. Furthermore, WormID.org provides links to the data corpus and related tools to work with whole-brain structural and activity images, convert datasets to NWB, and supplies tutorials and instructions for using these tools. Our aim is that this data standard, data corpus, and atlas of cell positions will be a continually evolving resource for the *C. elegans* neuroscience community, and eventually other model organisms.

### Analysis of biological factors in neuron positions

We statistically analyzed the spatial positions of *C. elegans* neuronal somas across individuals, strains, and lab conditions based on the mean and covariances in the statistical atlas. We focused on relative pairwise displacements rather than absolute positions because the absolute position of cells is dependent on positioning and deformation of the animal’s body during recording and thus requires global alignment of all animals. Aligning multiple animals into identical positions is an imperfect task. In contrast, measuring pairwise cell positions does not require alignment and is relatively robust against animal positioning and deformation.

Before analyzing statistical properties on the neuron positions, we assessed the percentage of neurons that were labeled by humans in each dataset. We found that neurons in the ventral ganglion and retrovesicular ganglion were less commonly labeled than neurons in other ganglia. As is shown here and previously in Yemini et al. 2021^6^, neurons in the ventral and retrovesicular ganglia exhibit high relative positional variability, which may explain why fewer of them were confidently labeled by researchers (**Fig. 4a**). For this reason, we explored several different factors hypothesized to contribute to the organization and variability of relative cell positions: i) gangliar boundaries (e.g., basal lamina and abutting tissue) which may restrict cell movement within the coelem, ii) synaptic connectivity which may impose energetic costs dependent on neuronal proximity, and iii) developmental-time and cell lineage effects whereby recently divided cells (i.e., sister cells) remain close together and more distant relatives (e.g., mother and grandmother cells) end up further apart.

**Figure 4.**
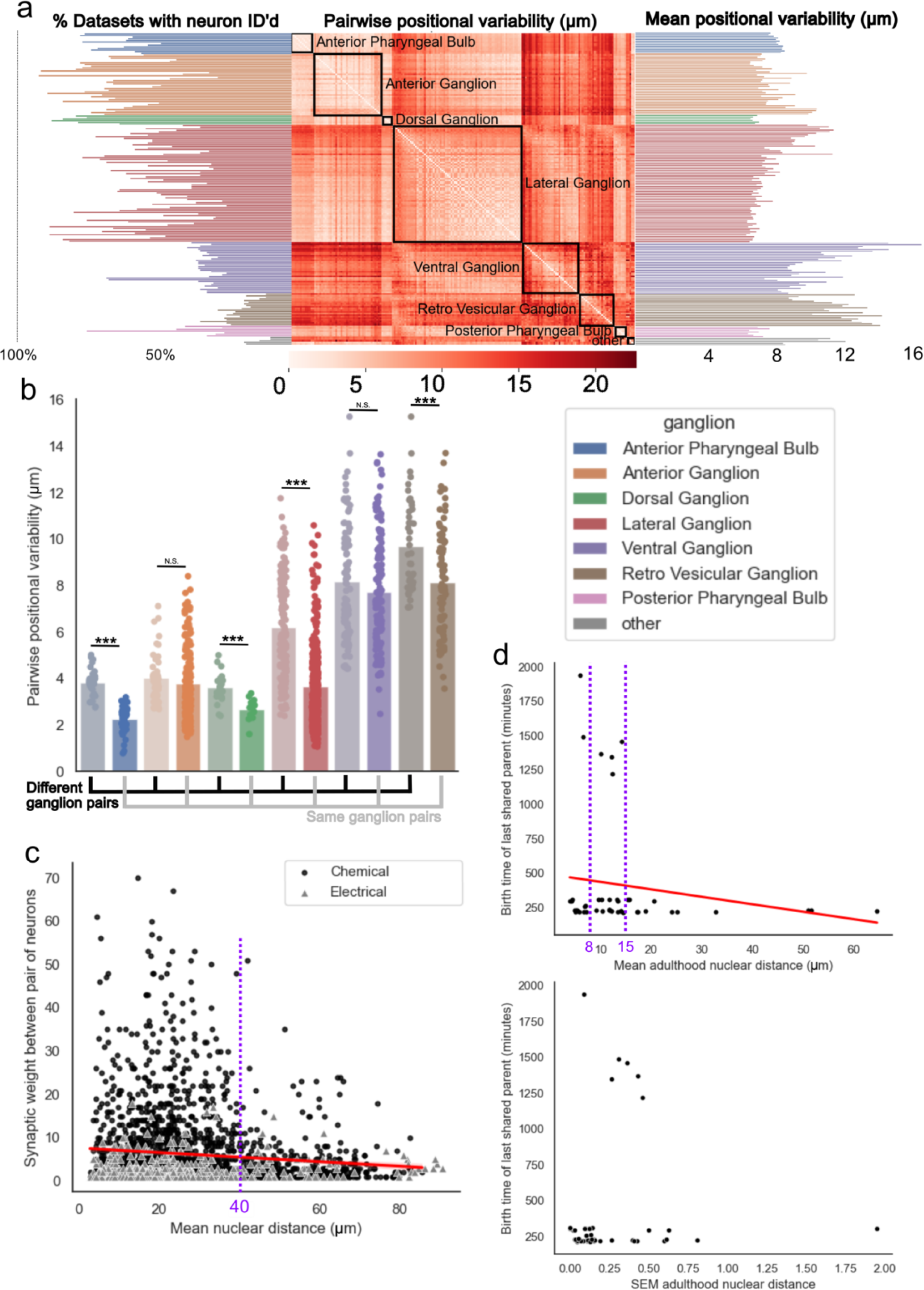
Analyses of neuron positions, distances, and positional variability. (a) Left: percentage of datasets containing each labeled neuron, organized anterior-posterior within each ganglion. Middle: heatmap of the STD of pairwise positional distances between each pair of neurons across datasets. Right: Averaged sums of heatmap rows. Neurons with higher mean positional variability have less stereotyped positions within the worm body. (b) Pairwise positional variability by ganglia for 10 closest neighbors of each neuron, separating neuron pairs in the same ganglion from pairs in different ganglia. Anterior pharynx: effect size 95% CI [-1.79, -1.38] μm, *p* = 2.5 ∗ 10^−21^, *N*_*same*_ = 53, *N*_*diff*_ = 36; Dorsal: effect size 95% CI [-1.16, -0.72] μm, *p* = 7.1 ∗ 10^−6^, *N*_*same*_ = 13, *N*_*diff*_ = 27; Lateral: effect size 95% CI [-2.80, -2.38] μm, *p* = 5.3 ∗ 10^−31^, *N*_*same*_ = 319, *N*_*diff*_ = 159; Retrovesicular: 95% CI effect size=[-1.97,-1.19]μm *p* = 9.3 ∗ 10^−5^, *N*_*same*_ = 89, *N*_*diff*_ = 43; Anterior: *p* = 0.161, *N*_*same*_ = 176, *N*_*diff*_ = 50; Ventral: *p* = 0.252, *N*_*same*_ = 52, *N*_*diff*_ = 37. (c) Relationship between pairwise neuron synaptic weights and their mean positional distance for chemical and electrical synapses. Chemical synapses: KendallTau τ = −0.036, *p* = 0.021, Pearson *R* = −0.098, *p* = 6.4 ∗ 10^−6^, N=2119; Electrical synapses: KendallTau τ = 0.009, *p* = 0.80, Pearson *R* = 0.031, *p* = 0.51, N=444. (d) Relationship between cell birth times and the mean and SEM of their nuclear positional distance in adulthood for sister cells. Mean: KendallTau τ = −0.144, *p* = 0.052, Pearson *R* = −0.161, *p* = 0.129; SEM: KendallTau τ = 0.014, *p* = 0.845, Pearson *R* = 0.074, *p* = 0.487.Most sisters are within 15 μm of each other in adulthood. More sisters that divide embryonically remain close together (<8 μm) than sisters that divide >16 hours later at postembryonic larval stages.

To test the first hypothesis, that gangliar boundaries regulate positional organization and variability, we measured the positional variability of neurons that are spatially close and compared pairs within the same ganglion to pairs straddling each other in different ganglia (see **Methods**). We found that neurons in the anterior pharyngeal bulb and neurons in the dorsal, lateral, and retrovesicular ganglia all exhibit significantly lower variability for pairs within the same ganglion compared to pairs in different ganglia. Conversely, for neurons in the anterior and ventral ganglia, we observed no significant difference between pairs in the same ganglion and pairs in different ganglia (**Fig. 4b**). We used an independent samples t-test to compare pairs within the same ganglion with those in different ganglia. Known anatomical features of the worm support this hypothesis: the pharynx is a muscular epithelial tube^47^ that rigidly encases neurons; the remaining ganglia are separated by basal lamina that loosely restricts their boundaries^44^; finally, the anterior and ventral ganglia (and comparatively smaller retrovesicular ganglion) are completely bounded whereas all other ganglia are open at least at one end, and in White et al. 1986 it had been noted that tight cellular packing in these regions led to “slop”, “uncertainty”, and in live animals even “flipping” from side to side of the cells contained therein. This finding suggests that neural identification algorithms could be improved by a hierarchical approach, such as first predicting ganglia membership, then predicting neuron identities within each ganglion.

Next, we explored the relationship between somatic distance and synaptic connectivity. Overall, there was a very weak but statistically significant correlation between nuclear distance and synaptic weight for chemical synapses and no significance or detectable correlation for electrical synapses. However, we found that nearby neurons (mean distance < 40 μm) exhibit a wide range of chemical synaptic weights ranging anywhere from 0 to 70 synapses (with a median synaptic count of 3), whereas distant neurons (mean distance > 40 μm) have a maximum synaptic count of ∼25 synapses (with a median count of 2) (**Fig. 4c**). This choice of distance cutoff was chosen by observing a distinct elbow at 40 μm in a 2D kernel density estimate plot of the scatter data (**Supplement 1)**. Our data suggests that neurons that are strongly wired together tend to be close to each other, although somatic proximity alone is not sufficient to imply strong connectivity. Recent findings in *C. elegans* have substantiated Peter’s rule: neurons with larger colocalized axodendritic regions are more likely to form connections.^48^ Our findings lend further support to this principle and suggest that close somatic or nuclear proximity also plays a role in determining neural connectivity.

Lastly, we explored the hypothesis that cell lineage is a determinant of adult cell positioning. Embryonic *C. elegans* are confined to a fixed volume within an eggshell approximately 50 μm in length and 30 μm in diameter.^49^ After hatching, they grow over 4x in length from birth (∼250 μm) to adulthood (over 1 mm), with an exponential expansion in their volume.^50,51^ Sister cells are cells whose lineage differs only at the very last division. We hypothesized that animal growth should lead to both larger distances and higher variability between older sister cells that divided in the embryo, versus younger sister cells born much later at postembryonic larval stages of development. Surprisingly, we found no statistically significant correlation between the time of cell division and nuclear distance or between time of cell division and distance variability measured by SEM (**Fig. 4d**). In fact, most sisters remained within 15 μm of each other (∼3 nuclei apart) at adulthood, regardless of when they were born. Strikingly, a substantial cohort of embryonic sisters ended up closer together at adulthood (< 8 μm) than those dividing at larval stages that occur more than 16 hours later (**Fig. 4d**). Our data rule out exponential postembryonic growth spurts as a major determinant of divergence and variability in neuron positions.

### Neuron identification performance increases for all laboratories and all tested algorithms when trained on a harmonized multi-lab corpus

Our previously published statistical atlas algorithm (“StatAtlas”) for automated neuron identification was trained on a homogenous dataset of 10 NeuroPAL worms.^6,28^ Formerly, this 10 worm training set achieved average accuracies of 86% overall in head neurons that ranged from 50% for the ventral ganglion to 100% for the anterior pharyngeal ganglion. These accuracies facilitate neural identification, but in practice they require substantial verification and manual corrections, and thus necessitating significant time expenditure in these tasks. Moreover, the algorithm fails to generalize to datasets produced by other labs (**Fig. 5b**). We tested the performance of our previously published algorithm on each of the six aggregated datasets. Initial performance on these datasets ranged from ∼21% to ∼65% with an average of 41%, (**Supplement 2**). This substantial decrease in accuracy on datasets from different labs exposes the limitation of using single-lab training sets to produce tools intended for use by different labs with different instrumentation, experimental methods, and data acquisition pipelines.

**Figure 5:**
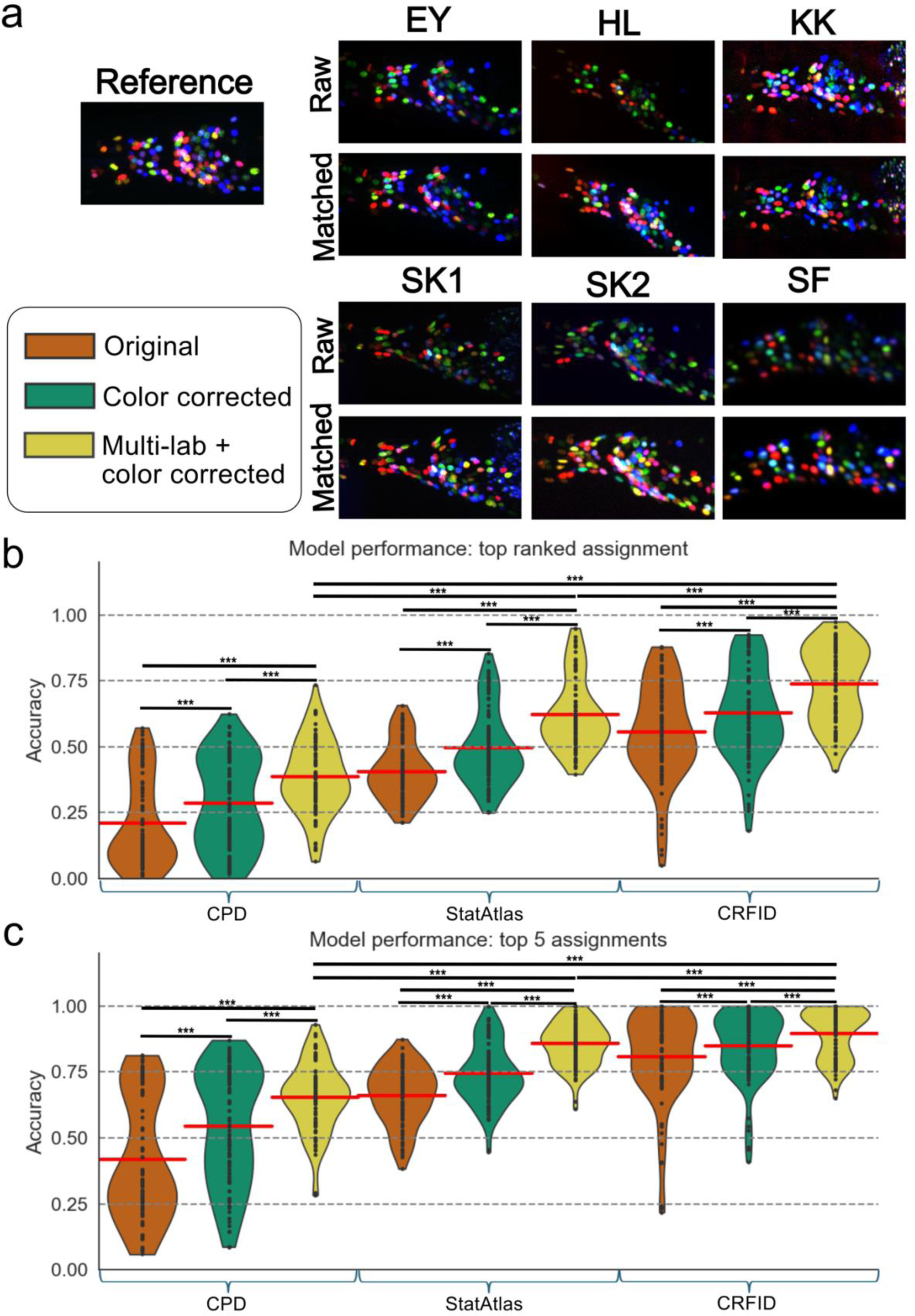
Improvements in neural identification accuracy. **(a)** Examples of raw and color-corrected (histogram matched) images from each lab and dataset. **(b)** Top ranked test accuracy for training set of original 10 reference worms with no color correction (orange), 10 reference worms with color correction (green), and the multi-lab corpus with color correction (yellow) for coherent point drift (CPD, left), the statistical atlas model (StatAtlas, middle), and the conditional random field model (CRF_ID, right). Algorithmic performance was evaluated using paired t-tests (where N=94 for each training set) to compare the performance of different atlases. Significance is reported using a Bonferroni correction with the convention of * for p<0.05, ** for p<0.01, and *** for p<0.001. **(c)** Same as b but using top 5 rank. Summary statistics and p values can be found in supplementary table 1.

To assess the performance benefits of using a large, harmonized corpus to train commonly-used automated neural-identification methods we tested two more popular algorithms: coherent point drift (“CPD”)^52^ and CRF_ID^26^. Coherent point drift is an untrained and unsupervised algorithm that: 1) globally aligns a sample point cloud of neurons to a reference atlas, then 2) locally matches points from the sample to their nearest neighbors in the atlas, and finally 3) identifies sample neurons (points) by their corresponding matches in the atlas. CRF_ID is a newer graph-based approach that identifies neurons using a combination of statistics from their individual features (e.g., absolute position and color) and pairwise relationships (e.g., displacement and angle relative to each other). There are no currently published benchmarks on the neural identification problem using CPD. Formerly, CRF_ID demonstrated a high accuracy of 83% when originally trained and tested solely on the HL dataset. Similar to StatAtlas, when testing the generalizability of the CPD and CRF_ID base models on the full WormID corpus we observed poor performance with an average overall accuracy of 39% and 59% respectively.

After inspecting recordings from multiple labs, we hypothesized that differences in color space may have negatively impacted algorithmic performance. Potential sources of color space variability include differences in microscope hardware, software and image settings, and configuration of the optical path. Anecdotally, in addition to these known sources of variability, researchers also typically adjust exposure, contrast, and other channel display parameters to make the composite rendered colors appear more like the images in the NeuroPAL reference manual.^53^ In aggregate, this suggested that harmonizing the color space may aid automatic algorithms.

We developed an approach to match the color histogram of a sample image to a reference histogram representing ideal coloring (see **Methods**). Histogram-matching the original small training set improved the accuracy of all three tested algorithms by an average of 8%, 9%, and 7% for CPD, statistical atlas, and CRF_ID respectively. It also qualitatively made composite color renderings better match the NeuroPAL reference manual, aiding users in annotating and correcting algorithmic predictions (**Fig. 5a,b,c**).

Given this success on the original small training set, we used the histogram-matched images to train a new atlas for the StatAtlas and CRF_ID algorithms on the full corpus of data. Test accuracy is reported using 5-fold cross-validation where each worm is tested against an atlas that was not trained on that worm. For CPD, we updated the algorithm to select the best template out of the full corpus (see **Methods** for further details). This led to significant improvement in accuracy across algorithms (**Fig. 5b, c)**, with an average improvement of 17%, 22%, and 18% that further raised average predictive accuracy from 22% to 39%, 41% to 62%, and 55% to 74% for CPD, StatAtlas, and CRF_ID respectively. This is equivalent to a ∼1.3x, ∼1.6x, and ∼1.7x reduction in error rate. Accuracy reached as high as 95% for several individual datasets for both StatAtlas and CRF_ID. Furthermore, when considering the top 5 neural identity assignments (rather than just the top 1), the multi-lab models showed average accuracies of 65%, 86%, and 89% for CPD, StatAtlas, and CRF_ID respectively, with some datasets reaching 100% accuracy for both StatAtlas and CRF_ID (**Fig. 5c, Supplement 3**). In addition, we see similar improvements in accuracy across most datasets for StatAtlas and CRFID when training using all except one dataset and then testing on the left-out dataset (**Supplement 6**). This indicates that most of the benefits from retraining come from achieving a better representation of the full diversity across datasets, rather than capturing the specific nuances of any one dataset. This generalizability will enable labs to use these retrained algorithms out of the box rather than needing to do additional fine-tuning on their own data.

Differences in accuracy between datasets may have been caused by a variety of factors including poor initial alignment, optical quality, non-neuronal artifacts in the images, and nonlinear deformations of the worm body. Additionally, datasets with less neurons annotated had better automatic labeling accuracy, presumably because experimenters only labeled the easiest neurons to identify and left the hardest ones unannotated (**Supplement 4,5**).

## Discussion

Aggregation and harmonization of data from a variety of different sources is necessary to build a corpus for analytical methods and machine learning tools that generalize across the diversity of real-world data. In this work, we present a data harmonization pipeline for analyzing whole-brain structural and activity imaging in *C. elegans*. This pipeline includes data aggregation, conversion to a standardized file format, software for analyzing these standardized datasets, pre-processing approaches to align images and color spaces, and spatial registration of sample neuron point clouds to a common atlas.

We used this corpus to study potential biological factors that organize cell position in *C. elegans.* Specifically, we find that: i) restrictions in bounding tissue and gangliar space likely contribute to variability in neuron positions, ii) neurons with somatic distances less than ∼40 μm of each other show higher synaptic connectivity, and iii) sister neurons that divide in the embryo can be found closer together at adulthood than ones dividing at larval stages more than 16 hours later. The positive relationship between synaptic connectivity and neuron somatic proximity thus augments the previously observed correlation of synaptic connectivity to axodendritic adjacency, termed Peter’s Rule. Moreover, the close distances and low positional variability we measured for embryonically born sister neurons rules out exponential organismal growth as a major cause in driving neurons apart from each other during the establishment of the adult Bauplan.

We then used the corpus to train a machine learning tool to automate the intensive task of labeling cells in these datasets. This substantially boosted generalized performance across datasets from contributing labs for each tested algorithm, despite the variability in data from these different groups. Accuracy of auto-identification now approaches human performance for certain datasets using the StatAtlas or CRF_ID algorithms. In the future, our corpus can be used to incorporate neuronal shape- and size-based descriptors as well as dynamical time-series features to further improve neural identification algorithms.

The WormID.org tools and resources are readily applicable to new whole-brain structural and activity imaging datasets, and these new datasets can be easily added to the existing corpus. These tools streamline public data sharing to facilitate both open science and to satisfy data-sharing mandates. We hope this resource will continue to grow in size and breadth to enable the development and benchmarking of new machine learning tools and algorithms. Our analyses of cell features based on the full corpus of data can immediately be used to inform better feature selection and algorithms that continue to improve automated approaches for neuron-subtype identification in volumetric images. Additionally, the large corpus and trained statistical atlas can serve as a descriptive resource of the underlying neurophysiology of *C. elegans*. Moreover, our resources can be incorporated into computational neurobiology courses, such as the Neuromatch Academy (neuromatch.io)^54^ to train the next global generation of neuroscientists on real-world datasets. As the community continues to develop new tools, this corpus will allow these new tools to be benchmarked for generalizable performance, spurring innovation.

As the community continues to scale up the generation of neural data and increasingly relies on machine learning analysis to tame this “big data”, there is an ever-growing need to unify disparate datasets to produce verifiably robust, accurate, and generalizable analytical approaches. Harmonization efforts such as ours can significantly reduce the activation energy necessary for collaboration, data sharing, and the development of unified community-wide tools across labs. While some of the resources we created are specific to *C. elegans,* the framework and much of our toolkit can be applied to other model organism imaging communities.

## Methods

### Standardized file format - Neurodata Without Borders

NWB is an HDF5-based format built specifically for neurophysiology data and has emerged as the de facto standard for storing neurophysiology datasets with associated metadata for reuse and sharing. NWB provides object types for data and metadata including acquisition parameters, segmentation of 3D image regions, fluorescent time series (e.g., for neural activity), experimental design information, multichannel electrophysiology time series data, 3D images, stimulus events during an experiment, and behavioral data.^42^

The base NWB schema supports two-dimensional structural and time-series multi-channel images but did not originally support the type of five-dimensional (multi-channel, volumetric, time-series) data that is used in *C. elegans* whole-brain activity imaging or other metadata associated with these types of experiments. To solve this problem, we developed ‘ndx-multichannel-volume’ as a novel extension to the existing Neurodata Without Borders (NWB) standardized file format. More information and resources about NWB can be found at nwb.org.

Our extension adds new objects built off of existing ones in the schema to add and improve support for multi-channel, volumetric, time-series images and the metadata associated with those images as well as volumetric segmentation data and metadata fields specific to *C. elegans* such as cultivation temperature and growth stage. This extension and the datasets presented in this work represent the first applications of the NWB data format to *C. elegans* and have now been incorporated as the standard for this model organism. This extension is flexible, open-source, and can be continuously updated to incorporate new types of data for future experiments.

### Storage on DANDI

Data and associated metadata were uploaded to the DANDI archive [RRID:SCR_017571] using the Python command line tool (https://doi.org/10.5281/zenodo.3692138). The data were first converted into the NWB format (https://doi.org/10.1101/2021.03.13.435173) and organized into a BIDS-like (https://doi.org/10.1038/sdata.2016.44) structure.

All datasets can be streamed or downloaded from the DANDI archive, available on WormID.org as well as these individual URLs:^55,56,57,58,59,60,61^

Original NeuroPAL: https://doi.org/10.48324/dandi.000715/0.240614.1942

EY: https://doi.org/10.48324/dandi.000472/0.240625.0454

HL: https://doi.org/10.48324/dandi.000714/0.240611.1954

KK: https://doi.org/10.48324/dandi.000692/0.240402.2118

SF: https://doi.org/10.48324/dandi.000776/0.240625.0015

SK1: https://doi.org/10.48324/dandi.000565/0.240625.0439

SK2: https://doi.org/10.48324/dandi.000472/0.240625.0450

### Software systems with embedded NWB I/O

We present two software examples with user-friendly GUIs to interface with NWB datasets and run standard analysis pipelines for cell segmentation, identification, tracking, and extracting time-series of neural-activity traces annotated with any experimental stimuli presented. First, we present the NeuroPAL_ID software (**Figure 2c**) for visualization, annotation, neuronal segmentation and identification, neural tracking, activity trace extraction and stimulus presentation data of volumetric NeuroPAL images and whole-brain activity. This software is pre-compiled for use on MacOS and Windows. The software is open-source, available from https://github.com/Yemini-Lab/NeuroPAL_ID/releases, and is written in MATLAB and Python. It has now been updated to include functionality described in this paper to enable histogram matching, color-corrected image visualization, and automated neural identification using the new Statistical Atlas. This software is written and managed by the Yemini Lab; further information can be found at https://www.yeminilab.com/neuropal.

Second, we present the eats-worm software for visualization, segmentation, and activity extraction of neural-activity time series from immobilized worms. Eats-worm similarly allows for manual verification and curation of the automatic segmentation and tracking algorithms. The tracking algorithm was optimized for tracking neurons across frames in immobilized worms, but there are currently efforts to extend this functionality to work for freely-moving worms as well. Eats-worm is written in Python and is built as a plugin to Napari, a popular 3D visualization tool. This software is written and managed by the Kato Lab; further information can be found at https://github.com/focolab/eats-worm.

Both software programs have embedded functionality to read and write NWB files. NWB I/O functionality enables a user to quickly run similar analyses on all of the datasets presented in this work without the need to develop specific pipelines to read in data from each dataset. Furthermore, this functionality can be easily embedded into MATLAB or Python-based analysis software.

### Data acquisition

NeuroPAL structural volumes and neural activity time series volumes were acquired using the protocols outlined in Yemini et al. 2021.^6^ After collection of these images, neurons were segmented and annotated according to the guidelines in the NeuroPAL manual.^53^ Specific immobilization methods, microscope setup, and experimental protocols differ slightly between datasets. All datasets were taken using spinning disk confocal microscopes with XY resolution varying from 0.1604-0.54 μm/pixel and Z resolutions varying from 0.54-1.5 μm/pixel. XY resolution was the same for NeuroPAL structural images and neural-activity images (using GCaMP6s) for all datasets, but Z resolution varied from 0.54-3 μm/pixel. Z resolution is generally lower for neural-activity images due to limitations in optical sectioning with confocal microscopes. Lower Z resolution also helps to reduce the number of frames needed to record a full volume for a single time-point, to aid imaging at a higher temporal resolution. Most images were taken with the worm immobilized in a microfluidic chip with the exception of the KK dataset (where worms were semi-restricted in a microfluidic device) and the SF dataset (where worms were freely moving). The NWB files and the DANDI datasets that hold them contain metadata for the specific setup and conditions in each dataset. For published datasets, additional information can be found in the associated publications.^6,7^

After acquisition of NeuroPAL structural volumes and whole-brain activity time-series, images were segmented using various automatic segmentation algorithms ranging from classical computer vision approaches (e.g., template matching^64^) to deep neural network approaches. These were then manually verified. Ground truth annotations were done using a combination of existing automatic identification algorithms followed by manual corrections. Each neuron identity label was either explicitly annotated by experts or manually verified after algorithmic identification. Note that varying levels of completeness in labeling are due to the difficulty of this manual annotation task. For several datasets with lower image quality, even experts could only confidently label 30-50% of segmented neurons in the volume. For neural activity time-series, neuron centers were first tracked across images using various algorithms and then manually verified by experts.^62,63^ Fluorescence activity is then extracted from these tracked ROIs to obtain time series of neural-activity traces. Neurons in the NeuroPAL structural volume were then matched to the ROIs in the neural activity time-series to get labeled activity traces.

Datasets from various labs were converted to the NWB standardized file format using the ndx-multichannel-volume extension presented in this work. These files were then uploaded to the DANDI archive where they are now publicly accessible for data streaming, download, or online visualization.

| Dataset | Microscope | Length of recording | Sample rate | Resolution (um/pixel) | Strain | Setup |
| --- | --- | --- | --- | --- | --- | --- |
| NP_og | Zeiss LSM880 spinning disk confocal | ~4 min | ~4Hz | 0.208 x 0.208 x 1.02 | OH16230 | Microfluidic chip |
| SF | Andor spinning disk confocal w/ Nikon ECLIPSE Ti microscope 40x water immersion | ~15 min | 1.7 Hz | 0.54 x 0.54 x 0.54 | Various | Freely moving |
| SK1 | Leica DMI8 inverted spinning disk confocal, 40x WI, 1.1 NA | ~25 min | 1.04 Hz | 0.1604 x 0.1604 x 1 (3 for calc images)<br>OR<br>0.3208 x 0.3208 x 0.75 (2.5 for calc images) | FC121, FC128, OH16230 | Microfluidic chip |
| SK2 | Leica DMI8 inverted spinning disk confocal, 40x WI, 1.1 NA | ~15 min | 3.3 Hz | 0.3208 x 0.3208 x 0.75 (1.5 for calc images) | OH16230 | Microfluidic chip |
| KK | Nikon Eclipse Ti-U inverted spinning disk confocal, 40x 1.3 NA | ~15-20 | 1.67 | 0.32 x 0.32 x 1.5 for both images | KDK92 | Semi-restricted in microfluidic device |
| HL | Perkin Elmer spinning disk confocal 1.3 NA, 40x oil OR Brucker Opterra II swept field confocal 0.75 NA, 40x air | NA | NA | 0.33 x 0.33 x 1 | OH15495 | Microfluidic device |
| EY | Spinning disk confocal | ~4 min | ~4 Hz | 0.27 x 0.27 x 1.5 | OH16230 |  |

### Butterfly plot

To produce the butterfly plot, we first manually found three orthogonal basis vectors to align neuron point clouds to a new cartesian coordinate space. To do so, we used human-guided affine transformation to roughly align these basis vectors to the anterior-posterior, dorsal-ventral, and left-right axes. The xyz coordinates of each neuron were projected into this new cartesian coordinate space and then converted to cylindrical coordinates by the following equations. We plotted the new *x* and θ coordinates on a 2D plane to get the butterfly plots shown in (**Fig. 1a**, **Fig. 5a**). This projection is akin to flattening the positions of the neurons along the circumference of a cylinder of the worm body and then unrolling that cylinder into a flattened plane.

### Histogram matching

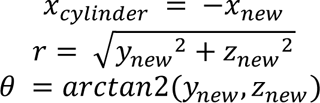

We modified the established approach of histogram matching to apply to 3-D volumetric, multi-channel data.^64^ We created a reference histogram using the 10 worms from the original NeuroPAL work. This data is stored as uint16, so there are 65,536 possible values for each pixel. For each channel, we created a histogram counting the number of pixels within the bin edges, assigning each color value its own bin, and then averaged the values in each of these bins across the 10 images. Practically, these histograms were very similar across these 10 datasets, so the averaged histogram looked similar to each of the individual histograms.

To color match a new animal sample, we calculated a histogram for each channel. The number of bins for each channel histogram was equal to the maximum intensity value present in that channel in the image. Practically, this means that there are a different number of histogram bins for each channel in each image because images were collected at different bit depths and with varying levels of saturation.

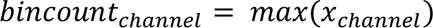

We then calculated a cumulative density at each color value for both the sample and the reference. We created a lookup table M to associate each gray count value x in the sample to the color value in the reference with the closest cumulative density. Next we created a new matched image with each pixel transformed into the new color space using this lookup table as shown below.

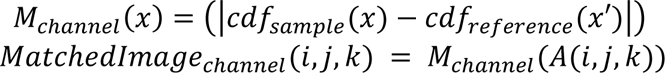

### Color extraction

To extract the color values for the neurons in each image, we first calculated the mean and standard deviation of the pixel gray counts in each channel, and then converted each pixel value into its Z-score based on its gray count value. We then took a sample of a 3×3×1 grid of pixel values around each segmented neuron center in each channel. We use the median values of this 3×3×1 grid as the RGB values for that neuron center. Color values were extracted post-histogram matching when training or testing using histogram-matched images. For non-histogram-matched images, there are no additional color pre-processing steps beyond Z-scoring.

### Positional variability analysis

We calculated pairwise positional variability by measuring the Euclidean distance between every pair of canonical head neurons across each structural volume when both neurons in that pair had a ground truth label. We then took the average and standard deviation of these distances for each neuron pair to find mean nuclear distance and pairwise positional variability, respectively. For these analyses we ignored pairs that are not present in at least 5 datasets. We used pairwise positional variability instead of absolute positional variability because absolute position is extremely sensitive to point-cloud realignment, which would make it hard to disaggregate natural positional variability from alignment errors; and furthermore, we are not interested in only how individual cells vary but rather how cells vary relative to each other. To get the mean positional variability for a given neuron, we averaged the mean pairwise distance for all pairs that contained that neuron.

Intra vs inter ganglion measures: for every neuron in the atlas, we found its n closest atlas neighbors and only measured pairs for its n closest neighbors. We then separated these pairings based on whether the two neurons in the pair are within the same ganglion or in two different ganglions. Note that pairs in different ganglions will appear twice: e.g. if one neuron in the pair is in the anterior ganglion and the other is in the lateral ganglion, the pair will be counted in the analysis for both the anterior ganglion and the lateral ganglion (**Fig. 4b**). Pairs within the same ganglion are only counted once. We compared this approach for n =1-20 (**Supplement 7**). For all numbers of neighbors there is higher positional variability for neighbors in different ganglia when compared to the same ganglion. The pattern stabilizes around n=7 and holds steady through n=20. Therefore, we selected 10 to use for n in our analysis.

Synaptic connection: synaptic weights between neuron pairs are derived from the whole-brain connectome of the adult hermaphrodite in Cook et al. 2019.^45^

Lineal distance: the cell lineage tree and associated birth times were taken from Sulston et al. 1982.^63^ The last shared parent cell between two neurons is the most recent shared parent node in the lineal tree. We used the birth time of the last shared parent cell between two neurons as the lineal distance and explored the relationship between this lineal distance and mean pairwise nuclear distance (**Fig. 4d**). In this analysis, we focus only on sister cells: terminal cells that only divided from each other at the very last stage of their lineal tree.

### Coherent Point Drift (CPD)

Coherent point drift has been a common algorithm for registering two similar point clouds to each other since its introduction in Myronenko and Song 2009^52^. CPD allows for both rigid and non-rigid point set registration. CPD models one point set as a set of GMM centroids that are fit to the second point set by maximizing the likelihood. GMM centroids are set to move coherently to preserve the structure of the point clouds. In the rigid case, the algorithm learns an affine transformation of the GMM centroid locations while in the non-rigid case, the algorithm learns a displacement function on the original centroid positions with an enforced regularization term to enforce smoothness. The objective function is optimized using an iterative EM optimization approach and yields both the aligned point set as well as an NxM correspondence probability matrix that represents the likelihood that each point n in set 1 corresponds to each point m in point set 2.

In this paper, we use the specific implementation of CPD used in Yu et al. 2021^34^ First, rigid CPD is used to roughly align a test worm point cloud to a template point cloud. Then, non-rigid CPD is used to model non-linear deformations between the semi-aligned test and template. Neuron assignments are then determined by creating a matrix of pairwise Euclidean distances between every neuron’s position and color in the test and every neuron in the template in the aligned space. We then use the Hungarian algorithm on this distance matrix to find the optimal label assignments. To get 2^nd^ ranked assignments, we assigned an infinite cost to each label assignment from the first pass and reran the Hungarian algorithm. We repeat this for the 3rd-5th order assignments.

Accuracy was calculated by counting the number of neurons whose algorithmic assignment was the same as the ground truth label and then dividing by the total number of neurons that have a ground truth label. Note that neurons without a ground truth label were not included in the accuracy metric but are still part of the cost matrix and received neuron assignments. Since there is no ground truth for these neurons we did not determine the accuracy of their label assignments.

The difference between the use of the ‘original (10) worms’ versus ‘multi-lab corpus’ for CPD is what set is included in possible options for the template. For the ‘original’ group, we compare every test set to each of the original 10 neuroPAL worms and report the accuracy for the template that has the highest average probability of correspondence after the rigid alignment step. Similarly, for the ‘multi-lab corpus’, each test worm is compared to each possible template worm in the whole multi-lab corpus and accuracy is reported similarly. The accuracy of CPD is highly sensitive to a good rough initial alignment and to similarity of the template and the test point cloud. The template with the highest average probability of correspondence is not necessarily the template that yields the highest accuracy, but it is the template that the algorithm has the highest confidence that it has found the ‘correct’ correspondence.

### Statistical Atlas training and inference

Statistical atlases used for testing performance were trained using the algorithm described in Varol et al. 2020.^28^ This algorithm uses a training set of neuron point clouds with both XYZ and RGB values and takes a block-coordinate descent approach where it iteratively learns affine transformation parameters to align the neuron point clouds, then updates the means and covariances of the positions and colors of each neuron until reaching convergence. This process generates mean and covariance parameters for each neuron as well as an aligned coordinate space for all the worms in the training set. The trained atlas consists of a list of neuron names alongside their associated means and covariances in the aligned position and color space.

We trained three atlases: the original atlas trained on just the original 10 NeuroPAL worms from Yemini et al. 2021^6^, the color corrected atlas trained on these same 10 worms after histogram matching, and the multi-lab + color-corrected atlas which is trained on the full corpus of histogram-matched data.

For the original atlas, we tested every dataset in the full corpus without histogram matching. For the color-corrected atlas, we similarly tested every dataset on the full corpus of data with histogram matching. For the atlas trained on the full corpus, we use K-fold cross-validation. The corpus was split into five equally-sized groups. For each group, an atlas was trained on all datasets in the other four groups and performance was reported for the out-of-training set group. The 10 worms used to train the original and color-corrected atlas were included in the training for each of these five groups. These 10 worms were not used to report testing accuracy for any of the atlases (**Fig. 5**). The fully trained atlas presented in **Fig. 3** was trained using the 10 original worms and the full corpus of data presented in this work, without splitting it into groups. This full atlas was embedded into the autoID functionality of the NeuroPAL ID software shown in **Fig. 2**.

Neuron point clouds used for testing were pre-aligned by learning an affine transformation from each sample dataset to the aligned coordinates of the atlas based on a subset of the ground truth labeled neurons in the sample. Briefly, assuming N neurons in the test sample and M neurons in the atlas, we calculated an NxM cost matrix using the Mahalanobis distance between each neuron center in the sample and each neuron distribution in the atlas. <u>*x*_*i*_</u> represents the XYZRGB values of neuron i, while <u>μ*x*_*j*_</u> and μ_*j*_ represent the XYZRGB mean and covariance respectively for neuron j in the atlas.

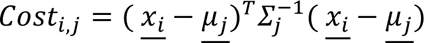

We then treated this cost matrix as a linear sum assignment problem. Label assignments (for neural identification) were calculated using the Hungarian algorithm.^43^ 2^nd^-5^th^ order ranked assignments and accuracy are calculated in the same way as described for CPD.

### CRF_ID training and inference

CRF_ID atlases and inference are conducted using the algorithm described in Chaudhary et al. 2021^26^. This approach follows a probabilistic-graphical-model framework based on conditional random fields. The graph is defined by node features corresponding to unary measures for each neuron center such as position and color and edge features corresponding to pairwise measures for each pair of neurons such as distance, relative angle, or probability that one neuron is anterior to the other. After features are selected, a data-driven atlas is trained on a corpus of data to determine the average values for each of the measured features; then for a test worm, node and edge potentials are calculated based on comparison of each feature in the test worm to the atlas and infer the best global assignment of labels by maximizing an energy function using an approximate inference method. For the analysis in this work, we used the color information solely to define the node potentials, and the pairwise angle relationships only to define the edge potentials. Optimizing the weights of the node and edge features may result in a higher prediction accuracy.

We trained three atlases: the original atlas trained on just the original 10 NeuroPAL worms from Yemini et al. 2021^6^, the color corrected atlas trained on these same 10 worms after histogram matching, and the multi-lab + color-corrected atlas which is trained on the full corpus of histogram-matched data. This training approach follows the same K-fold cross validation approach used for the Statistical Atlas method.

We use the roughly pre-aligned point clouds used in the Statistical Atlas algorithm as input to the CRF_ID algorithm to eliminate possible difference in the initial alignment step, which can dramatically change accuracy.

In practice, there are nearly always fewer detected neuron centers in a given image than total cells in the atlas. CRF_ID handles this by modeling a hidden variable ***h*** ∈ {0,1}^*N*^ where N is the number of neurons in the atlas. This variable specifies the probability that a given cell is missing in the image. Based on the number of cells in the test image, P cells are uniformly selected across different regions of the head and removed from the atlas. This process is repeated ∼1000 times to sample multiple possible combinations of **h.** The top 1-5 predicted assignments are generated by compiling a list of the most frequent labels for each cell in the test image across all runs. Accuracy is reported in the same way as CPD and CRF_ID.

Optimizing the aforementioned energy function using an approximate inference method produces marginal distributions of label assignments for each cell. The top 1-5 predicted label assignments for each cell were generated by sorting the marginal probability of labels in a descending order. The label that resulted in the highest marginal probability was assigned as top 1.

## Author contributions

D.S. developed the NWB extension format, harmonization algorithms and code, and performed data analysis. K.R. updated the NeuroPAL ID software and performed data analysis. J.B., S.B. and R.D. developed the eats-worm software. H.J.L. and Z.Y. performed data analysis. D.S, K.R., G.B., R.D., J.B., G.C., C.W., R.S., S.C., H.L., and E.Y. performed collection, curation, and annotation of included datasets. C.W., R.S., S.O., and K.K. contributed to data conversion and the NWB extension format. B.D. and R.L. contributed to the NWB extension format. G.B. developed visualization approaches. D.S., E.Y. and S.K. designed the analysis, interpreted data, and wrote the manuscript. All authors reviewed the manuscript.

## Acknowledgements

We thank Erdem Varol and Amin Nejatbakhsh for providing code and troubleshooting support to train the statistical atlas, and to label images of dense neurons used in figure panels that show atlases of neuron positions. We thank Adam Atanas and Steve Flavell for contributing their datasets. Research funding was provided by National Institute of General Medical Sciences of the National Institutes of Health 582 (#R35GM124735, SK), the Weill Institute for Neurosciences, and the Weill Neurohub. Development of eats-worm is supported by a napari Plugin Foundation grant from the Chan Zuckerberg Initiative. EY was funded by the Esther A. & Joseph Klingenstein Fund, the Simons Foundation, and the Hypothesis Fund.

## Supplemental Information

**Supplement 1:**
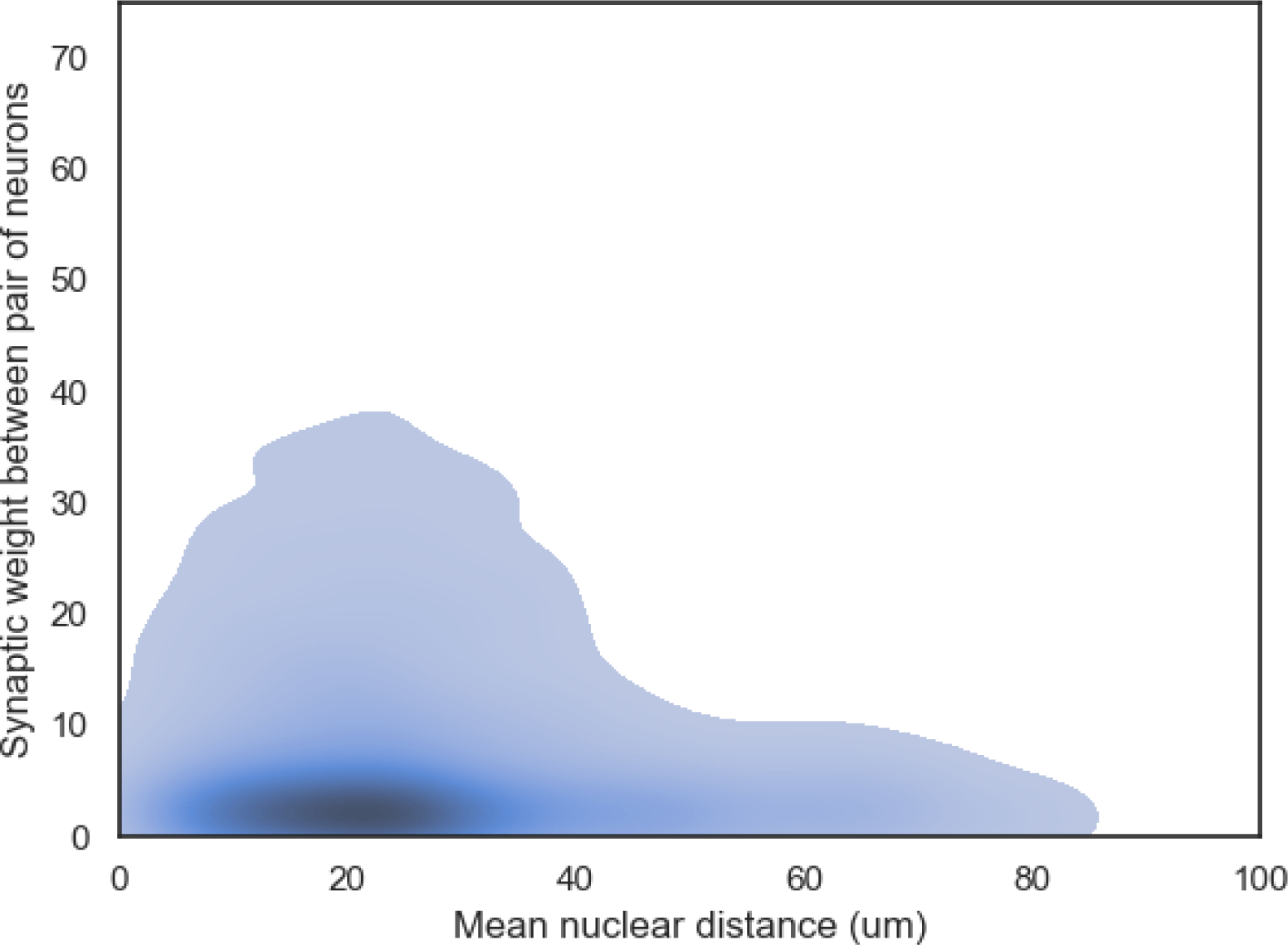
KDE plot of mean nuclear distance and synaptic weight between pairs of neurons. Contour encompasses 95% of the cumulative mass of the data. Color shows density with higher density colored darker.

**Supplementary table 1:** Summary statistics for figure 5.

| N=94 | Top1 mean effect size<br>[95% CI] | Top1 p-value | Top5 mean effect size<br>[95% CI] | Top5 p-value |
| --- | --- | --- | --- | --- |
| CPD: Original vs color corrected | 7.66% [5.07%, 10.25%] | 6.298e-08 | 12.63% [9.41%, 15.85%] | 8.959e-12 |
| CPD: Original vs multi-lab + color corrected | 17.62% [13.60%, 21.63%] | 9.881e-14 | 23.44% [18.71%, 28.17%] | 4.119e-16 |
| CPD: Color corrected vs. multi-lab + color corrected | 9.96% [6.69%, 13.23%] | 2.958e-08 | 10.81% [7.20%, 14.42%] | 4.398e-08 |
| StatAtlas: Original vs color corrected | 9.05% [7.60%, 10.50%] | 2.217e-21 | 8.30% [7.12%, 9.47%] | 1.117e-24 |
| StatAtlas: Original vs multi-lab + color corrected | 21.57% [19.71%, 23.42%] | 2.373e-40 | 19.76% [17.51%, 22.00%] | 3.449e-31 |
| StatAtlas: Color corrected vs. multi-lab + color corrected | 12.52% [11.18%, 13.86%] | 4.698e-33 | 11.46% [9.68%, 13.24%] | 3.172e-22 |
| CRF_ID: Original vs color corrected | 7.30% [5.09%, 9.51%] | 2.829e-09 | 4.11% [2.19%, 6.02%] | 4.948e-05 |
| CRF_ID: Original vs multi-lab + color corrected | 18.39% [15.16%, 21.62%] | 3.222e-19 | 8.70% [5.53%, 11.86%] | 3.839e-07 |
| CRF_ID: Color corrected vs multi-lab + color corrected | 11.09% [8.44%, 13.75%] | 7.682e-13 | 4.59% [2.52%, 6.67%] | 2.910e-05 |
| Multi-lab + color corrected: CPD vs StatAtlas | 23.61% [20.82%, 26.40%] | 6.064e-30 | 20.51% [18.17%, 22.85%] | 4.519e-31 |
| Multi-lab + color corrected: CPD vs CRF_ID | 35.24% [31.80%, 38.67%] | 4.167e-36 | 24.17% [21.26%, 27.07%] | 1.825e-29 |
| Multi-lab + color corrected: StatAtlas vs CRF_ID | 11.63% [9.09%, 14.16%] | 1.492e-14 | 3.66% [2.52%, 4.79%] | 5.908e-09 |

**Supplement 2:**
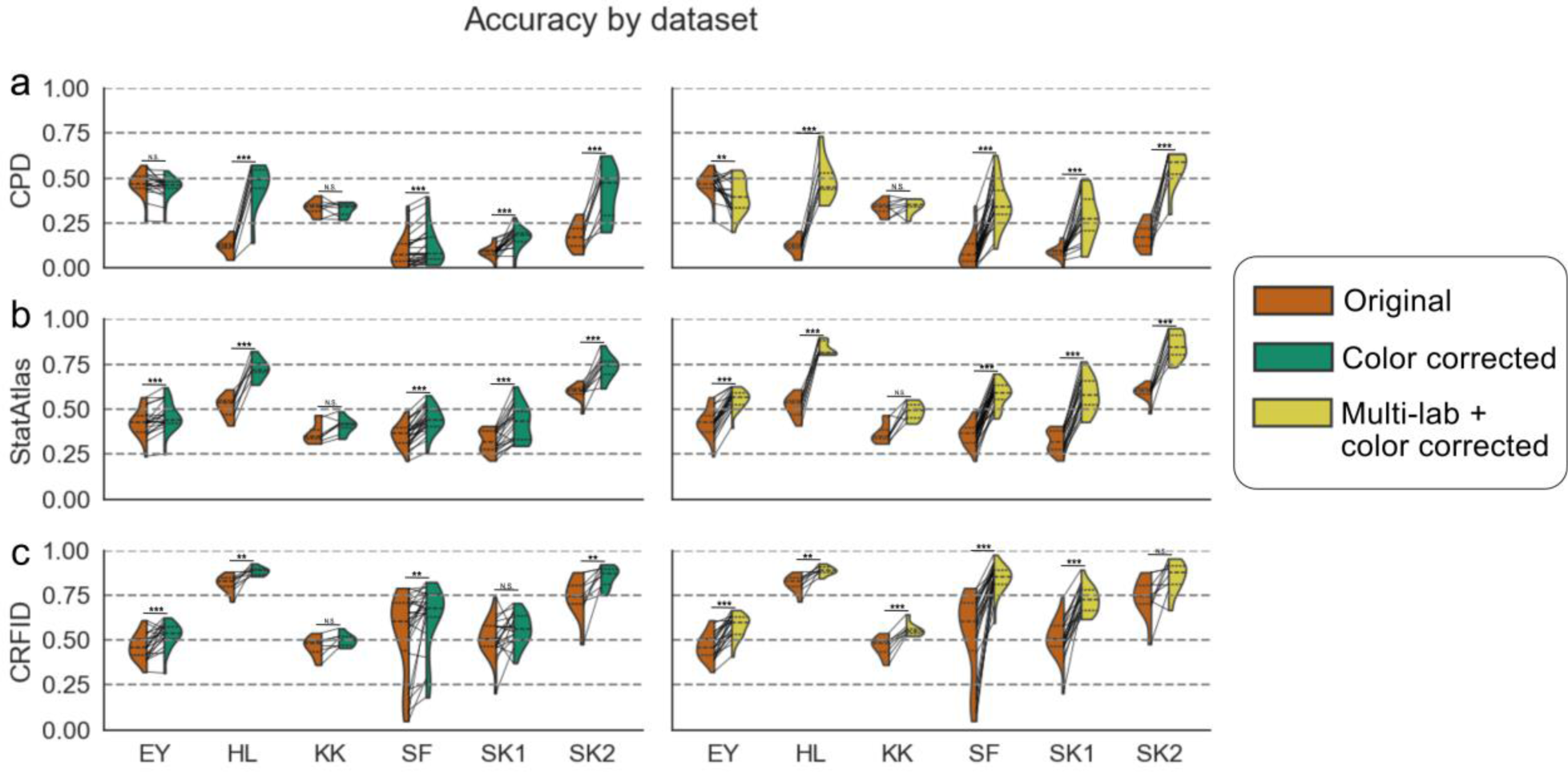
Top accuracy by dataset **(a, b, c)** Top ranked test accuracy for training set of original 10 reference worms with no color correction (orange), 10 reference worms with color correction (green), and the multi-lab corpus with color correction (yellow) broken out by model. Algorithmic performance was evaluated using paired t-tests (where N=94 for each training set) to compare the performance of different atlases. Significance is reported using a Bonferroni correction with the convention of * for p<0.05, ** for p<0.01, and *** for p<0.001. **(a)** Accuracy by dataset for CPD. EY dataset: original (10 worms) no color correction versus with color correction - mean effect size = -2.00%, 95% CI effect size = [-3.60%, -0.39%], p = 2.491e-02; original with no color correction versus multi-lab with color correction - mean effect size = -6.59%, 95% CI effect size = [-10.85%, -2.34%], p = 6.695e-03. HL dataset: original (10 worms) no color correction versus with color correction - mean effect size = 33.75%, 95% CI effect size = [27.57%, 39.93%], p = 6.815e-06; original with no color correction versus multi-lab with color correction - mean effect size = 37.66%, 95% CI effect size = [29.39%, 45.94%], p = 2.624e-05. KK dataset: original (10 worms) no color correction versus with color correction - mean effect size = -0.67%, 95% CI effect size = [-3.01%, 1.67%], p = 6.075e-01; original with no color correction versus multi-lab with color correction - mean effect size = 1.42%, 95% CI effect size = [-1.58%, 4.43%], p = 4.041e-01. SF dataset: original (10 worms) no color correction versus with color correction - mean effect size = 2.96%, 95% CI effect size = [1.83%, 4.09%], p = 2.671e-05; original with no color correction versus multi-lab with color correction - mean effect size = 26.09%, 95% CI effect size = [21.17%, 31.02%], p = 1.469e-10. SK1 dataset: original (10 worms) no color correction versus with color correction - mean effect size = 7.39%, 95% CI effect size = [5.08%, 9.70%], p = 5.516e-06; original with no color correction versus multi-lab with color correction - mean effect size = 19.55%, 95% CI effect size = [14.16%, 24.95%], p = 1.010e-06. SK2 dataset: original (10 worms) no color correction versus with color correction - mean effect size = 23.88%, 95% CI effect size = [17.33%, 30.44%], p = 6.968e-05; original with no color correction versus multi-lab with color correction - mean effect size = 37.46%, 95% CI effect size = [30.69%, 44.24%], p = 2.406e-06. **(b)** Accuracy by dataset for StatAtlas. EY dataset: original (10 worms) no color correction versus with color correction - mean effect size = 3.43%, 95% CI effect size = [1.76%, 5.10%], p = 6.888e-04; original with no color correction versus multi-lab with color correction - mean effect size = 12.85%, 95% CI effect size = [10.79%, 14.91%], p = 1.080e-10. HL dataset: original (10 worms) no color correction versus with color correction - mean effect size = 20.67%, 95% CI effect size = [16.96%, 24.38%], p = 5.871e-06; original with no color correction versus multi-lab with color correction - mean effect size = 31.95%, 95% CI effect size = [27.70%, 36.20%], p = 5.955e-07. KK dataset: original (10 worms) no color correction versus with color correction - mean effect size = 4.01%, 95% CI effect size = [1.70%, 6.31%], p = 1.403e-02; original with no color correction versus multi-lab with color correction - mean effect size = 12.22%, 95% CI effect size = [7.57%, 16.87%], p = 1.719e-03. SF dataset: original (10 worms) no color correction versus with color correction - mean effect size = 8.56%, 95% CI effect size = [7.01%, 10.11%], p = 6.380e-11; original with no color correction versus multi-lab with color correction - mean effect size = 22.85%, 95% CI effect size = [20.21%, 25.50%], p = 3.298e-15. SK1 dataset: original (10 worms) no color correction versus with color correction - mean effect size = 9.78%, 95% CI effect size = [6.80%, 12.75%], p = 3.856e-06; original with no color correction versus multi-lab with color correction - mean effect size = 25.88%, 95% CI effect size = [22.85%, 28.90%], p = 8.466e-13. SK2 dataset: original (10 worms) no color correction versus with color correction - mean effect size = 14.23%, 95% CI effect size = [10.56%, 17.91%], p = 4.361e-05; original with no color correction versus multi-lab with color correction - mean effect size = 26.06%, 95% CI effect size = [21.88%, 30.25%], p = 8.712e-07. **(c)** Accuracy by dataset for CRF_ID. EY dataset: original (10 worms) no color correction versus with color correction - mean effect size = 6.04%,95% CI effect size = [3.22%, 8.86%], p = 4.601e-04; original with no color correction versus multi-lab with color correction - mean effect size = 10.97%,95% CI effect size = [8.32%, 13.62%], p = 9.944e-08. HL dataset: original (10 worms) no color correction versus with color correction - mean effect size = 6.94%,95% CI effect size = [3.56%, 10.32%], p = 4.708e-03; original with no color correction versus multi-lab with color correction - mean effect size = 7.28%,95% CI effect size = [4.05%, 10.50%], p = 2.776e-03. KK dataset: original (10 worms) no color correction versus with color correction - mean effect size = 3.61%,95% CI effect size = [1.13%, 6.10%], p = 2.958e-02; original with no color correction versus multi-lab with color correction - mean effect size = 9.42%,95% CI effect size = [6.72%, 12.12%], p = 3.264e-04. SF dataset: original (10 worms) no color correction versus with color correction - mean effect size = 9.70%,95% CI effect size = [3.94%, 15.46%], p = 2.897e-03; original with no color correction versus multi-lab with color correction - mean effect size = 31.28%,95% CI effect size = [23.27%, 39.28%], p = 5.119e-08. SK1 dataset: original (10 worms) no color correction versus with color correction - mean effect size = 5.15%,95% CI effect size = [0.17%, 10.13%], p = 5.838e-02; original with no color correction versus multi-lab with color correction - mean effect size = 21.75%,95% CI effect size = [17.54%, 25.97%], p = 4.768e-09. SK2 dataset: original (10 worms) no color correction versus with color correction - mean effect size = 11.27%,95% CI effect size = [5.15%, 17.38%], p = 6.775e-03; original with no color correction versus multi-lab with color correction - mean effect size = 10.91%,95% CI effect size = [3.73%, 18.08%], p = 1.803e-02.

**Supplement 3:**
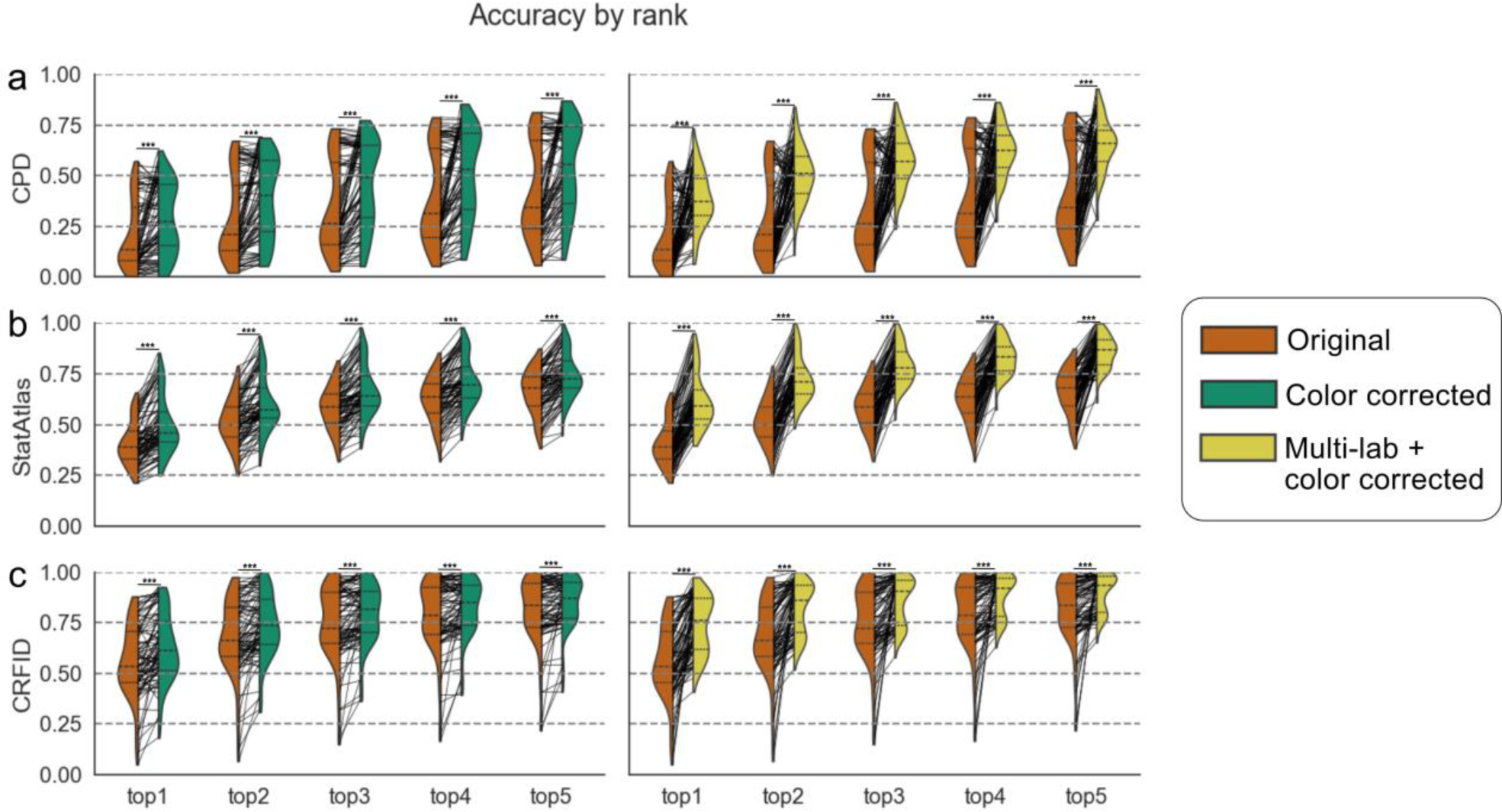
Top n ranked assignment accuracy broken out by training group and model. Algorithmic performance was evaluated using paired t-tests (where N=94 for each training set) to compare the performance of different atlases. Significance is reported using a Bonferroni correction with the convention of * for p<0.05, ** for p<0.01, and *** for p<0.001. **(a)** Accuracy by rank for CPD. Original (10 worms) no color correction versus with color correction: top1 - mean effect size = 7.66%, 95% CI effect size = [5.07%, 10.25%], p = 6.298e-08; top2 - mean effect size = 10.28%, 95% CI effect size = [7.42%, 13.14%], p = 2.018e-10; top3 - mean effect size = 11.52%, 95% CI effect size = [8.55%, 14.49%], p = 1.252e-11; top4 - mean effect size = 12.34%, 95% CI effect size = [9.16%, 15.52%], p = 1.326e-11; top5 - mean effect size = 12.63%, 95% CI effect size = [9.41%, 15.85%], p = 8.959e-12. Original with no color correction versus multi-lab with color correction: top1 - mean effect size = 17.62%, 95% CI effect size = [13.60%, 21.63%], p = 9.881e-14; top2 - mean effect size = 21.80%, 95% CI effect size = [17.31%, 26.28%], p = 1.047e-15; top3 - mean effect size = 22.64%, 95% CI effect size = [18.03%, 27.25%], p = 6.539e-16; top4 - mean effect size = 23.21%, 95% CI effect size = [18.50%, 27.91%], p = 5.260e-16; top5 - mean effect size = 23.44%, 95% CI effect size = [18.71%, 28.17%], p = 4.119e-16 **. (b)** Accuracy by rank for StatAtlas. Original (10 worms) no color correction versus with color correction: top1 - mean effect size = 9.05%, 95% CI effect size = [7.60%, 10.50%], p = 2.217e-21; top2 - mean effect size = 9.01%, 95% CI effect size = [7.62%, 10.40%], p = 1.964e-22; top3 - mean effect size = 8.66%, 95% CI effect size = [7.44%, 9.89%], p = 8.658e-25; top4 - mean effect size = 8.83%, 95% CI effect size = [7.60%, 10.06%], p = 3.591e-25; top5 - mean effect size = 8.30%, 95% CI effect size = [7.12%, 9.47%], p = 1.117e-24. Original with no color correction versus multi-lab with color correction: top1 - mean effect size = 21.57%, 95% CI effect size = [19.71%, 23.42%], p = 2.373e-40; top2 - mean effect size = 21.37%, 95% CI effect size = [19.31%, 23.42%], p = 1.468e-36; top3 - mean effect size = 20.91%, 95% CI effect size = [18.98%, 22.85%], p = 6.920e-38; top4 - mean effect size = 20.84%, 95% CI effect size = [18.71%, 22.96%], p = 1.310e-34; top5 - mean effect size = 19.76%, 95% CI effect size = [17.51%, 22.00%], p = 3.449e-31. (**c)** Accuracy by rank for CRF ID. Original (10 worms) no color correction versus with color correction: top1 - mean effect size = 7.30%, 95% CI effect size = [5.09%, 9.51%], p = 2.829e-09; top2 - mean effect size = 6.45%, 95% CI effect size = [4.30%, 8.60%], p = 4.148e-08; top3 - mean effect size = 4.98%, 95% CI effect size = [3.00%, 6.96%], p = 2.710e-06; top4 - mean effect size = 4.38%, 95% CI effect size = [2.42%, 6.34%], p = 2.473e-05; top5 - mean effect size = 4.11%, 95% CI effect size = [2.19%, 6.02%], p = 4.948e-05. Original with no color correction versus multi-lab with color correction: top1 - mean effect size = 18.39%, 95% CI effect size = [15.16%, 21.62%], p = 3.222e-19; top2 - mean effect size = 14.97%, 95% CI effect size = [11.47%, 18.47%], p = 2.851e-13; top3 - mean effect size = 11.55%, 95% CI effect size = [8.16%, 14.94%], p = 1.101e-09; top4 - mean effect size = 9.89%, 95% CI effect size = [6.58%, 13.19%], p = 4.730e-08; top5 - mean effect size = 8.70%, 95% CI effect size = [5.53%, 11.86%], p = 3.839e-07.

**Supplement 4:**
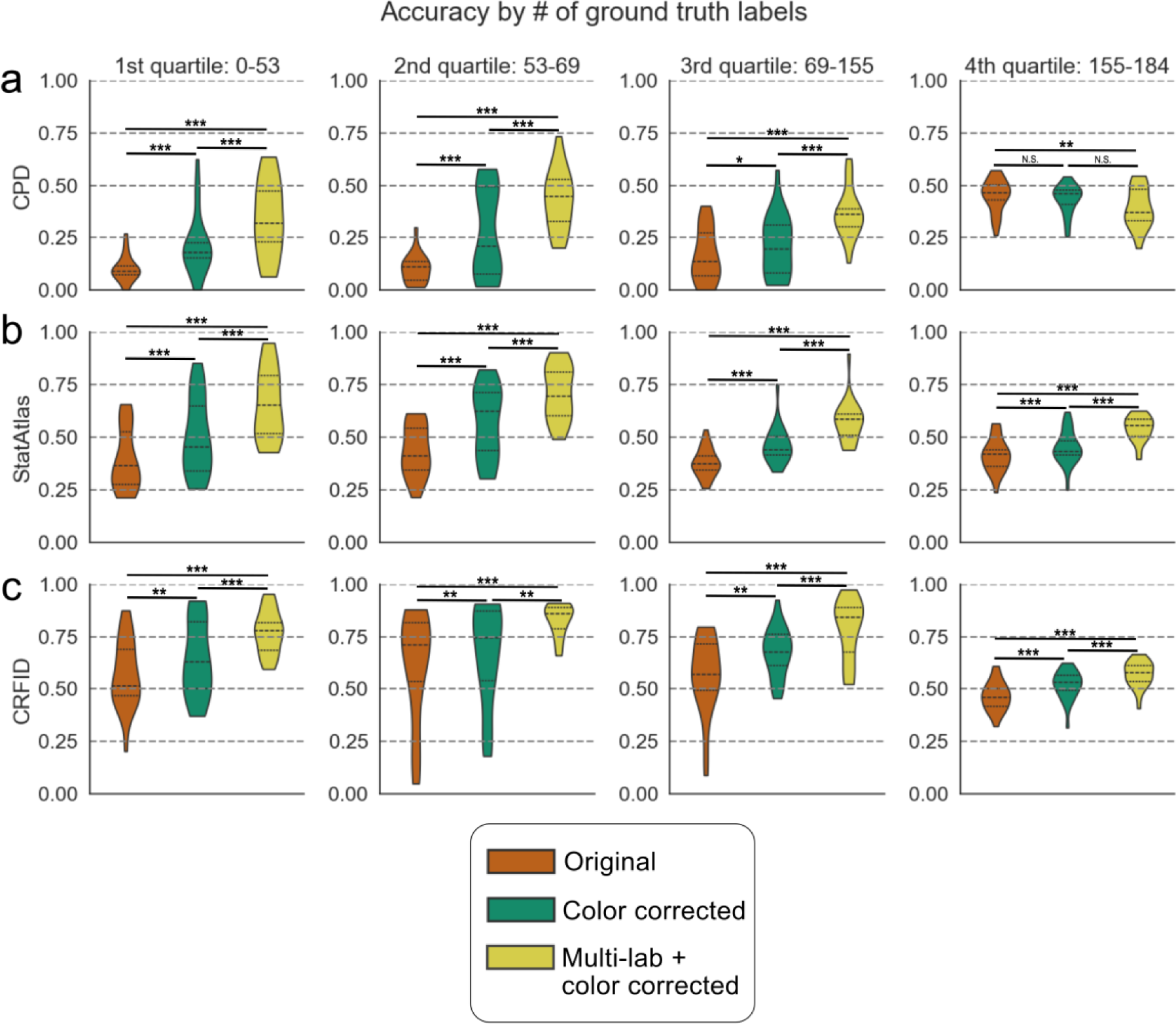
Top ranked neural identification accuracy for datasets with varying numbers of ground truth labels. All statistics are conducted with a paired t-test. First quartile N=25, second quartile N=26, third quartile N=27, fourth quartile N=16. Quartiles are determined by the number of ground truth labels in each dataset. Significance is reported using a Bonferroni correction with the convention of * for p<0.05, ** for p<0.01, and *** for p<0.001. **(a)** Accuracy by quartile for CPD. 1st quartile: original (10 worms) no color correction versus with color correction - mean effect size = 11.11%, 95% CI effect size = [7.25%, 14.96%], p = 1.162e-05; original with no color correction versus multi-lab with color correction - mean effect size = 24.40%, 95% CI effect size = [18.20%, 30.61%], p = 1.124e-07; original with color correction versus multi-lab with color correction - mean effect size = 13.30%, 95% CI effect size = [7.85%, 18.74%], p = 9.123e-05. 2nd quartile: original (10 worms) no color correction versus with color correction - mean effect size = 17.21%, 95% CI effect size = [10.39%, 24.03%], p = 6.144e-05; original with no color correction versus multi-lab with color correction - mean effect size = 32.96%, 95% CI effect size = [27.42%, 38.49%], p = 7.032e- 11; original with color correction versus multi-lab with color correction - mean effect size = 15.74%, 95% CI effect size = [8.48%, 23.00%], p = 3.335e-04. 3rd quartile: original (10 worms) no color correction versus with color correction - mean effect size = 4.58%, 95% CI effect size = [1.20%, 7.96%], p = 1.414e-02; original with no color correction versus multi-lab with color correction - mean effect size = 19.99%, 95% CI effect size = [14.47%, 25.51%], p = 3.175e-07; original with color correction versus multi-lab with color correction - mean effect size = 15.40%, 95% CI effect size = [9.73%, 21.08%], p = 2.166e-05. 4th quartile: original (10 worms) no color correction versus with color correction - mean effect size = -1.72%, 95% CI effect size = [- 3.26%, -0.19%], p = 3.829e-02; original with no color correction versus multi-lab with color correction - mean effect size = - 5.96%, 95% CI effect size = [-9.80%, -2.12%], p = 5.834e-03; original with color correction versus multi-lab with color correction - mean effect size = -4.23%, 95% CI effect size = [-7.51%, -0.96%], p = 1.864e-02. **(b)** Accuracy by quartile for StatAtlas. 1st quartile: original (10 worms) no color correction versus with color correction - mean effect size = 11.01%, 95% CI effect size = [8.35%, 13.68%], p = 5.059e-08; original with no color correction versus multi-lab with color correction - mean effect size = 26.44%, 95% CI effect size = [23.56%, 29.32%], p = 1.265e-14; original with color correction versus multi-lab with color correction - mean effect size = 15.43%, 95% CI effect size = [12.38%, 18.47%], p = 1.443e-09. 2nd quartile: original (10 worms) no color correction versus with color correction - mean effect size = 13.54%, 95% CI effect size = [10.27%, 16.81%], p = 4.780e- 08; original with no color correction versus multi-lab with color correction - mean effect size = 27.71%, 95% CI effect size = [25.22%, 30.20%], p = 2.244e-16; original with color correction versus multi-lab with color correction - mean effect size = 14.16%, 95% CI effect size = [11.53%, 16.80%], p = 4.772e-10. 3rd quartile: original (10 worms) no color correction versus with color correction - mean effect size = 8.22%, 95% CI effect size = [5.98%, 10.47%], p = 2.624e-07; original with no color correction versus multi-lab with color correction - mean effect size = 19.75%, 95% CI effect size = [16.14%, 23.36%], p = 2.053e- 10; original with color correction versus multi-lab with color correction - mean effect size = 11.52%, 95% CI effect size = [9.18%, 13.87%], p = 1.536e-09. 4th quartile: original (10 worms) no color correction versus with color correction - mean effect size = 3.68%, 95% CI effect size = [2.18%, 5.19%], p = 7.996e-05; original with no color correction versus multi-lab with color correction - mean effect size = 12.84%, 95% CI effect size = [11.00%, 14.68%], p = 1.557e-12; original with color correction versus multi-lab with color correction - mean effect size = 9.16%, 95% CI effect size = [7.32%, 10.99%], p = 1.189e-09. **(c)** Accuracy by quartile for CRF_ID. 1st quartile: original (10 worms) no color correction versus with color correction - mean effect size = 7.22%, 95% CI effect size = [2.25%, 12.18%], p = 9.428e-03; original with no color correction versus multi-lab with color correction - mean effect size = 19.97%, 95% CI effect size = [15.62%, 24.32%], p = 8.294e-09; original with color correction versus multi-lab with color correction - mean effect size = 12.75%, 95% CI effect size = [8.00%, 17.51%], p = 2.936e-05. 2nd quartile: original (10 worms) no color correction versus with color correction - mean effect size = 5.53%, 95% CI effect size = [2.51%, 8.55%], p = 1.646e-03; original with no color correction versus multi-lab with color correction - mean effect size = 21.42%, 95% CI effect size = [11.84%, 31.00%], p = 2.412e-04; original with color correction versus multi-lab with color correction - mean effect size = 15.89%, 95% CI effect size = [7.27%, 24.50%], p = 1.563e-03. 3rd quartile: original (10 worms) no color correction versus with color correction - mean effect size = 10.91%, 95% CI effect size = [5.08%, 16.75%], p = 1.304e-03; original with no color correction versus multi-lab with color correction - mean effect size = 21.73%, 95% CI effect size = [15.50%, 27.96%], p = 5.751e-07; original with color correction versus multi-lab with color correction - mean effect size = 10.82%, 95% CI effect size = [8.39%, 13.25%], p = 9.590e-09. 4th quartile: original (10 worms) no color correction versus with color correction - mean effect size = 5.45%, 95% CI effect size = [2.90%, 8.00%], p = 3.574e-04; original with no color correction versus multi-lab with color correction - mean effect size = 10.63%, 95% CI effect size = [8.26%, 13.00%], p = 8.450e-09; original with color correction versus multi-lab with color correction - mean effect size = 5.18%, 95% CI effect size = [3.34%, 7.02%], p = 1.358e-05.

**Supplement 5:**
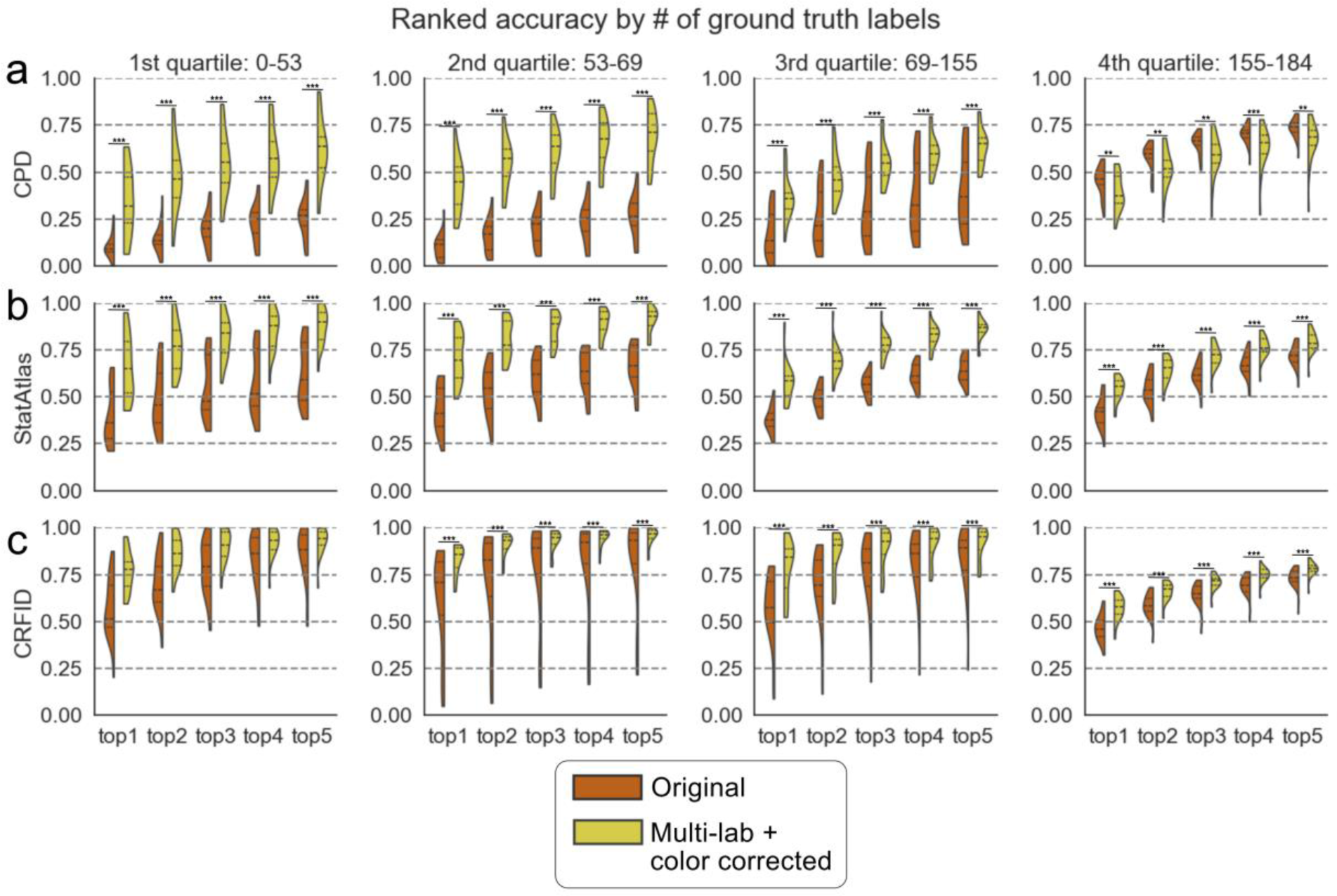
Top n ranked neural identification accuracy for datasets with varying numbers of ground truth labels. All statistics are conducted with a paired t-test. First quartile N=25, second quartile N=26, third quartile N=27, fourth quartile N=16. Quartiles are determined by the number of ground truth labels in each dataset. Significance is reported using a Bonferroni correction with the convention of * for p<0.05, ** for p<0.01, and *** for p<0.001. **(a)** Accuracy by rank and quartile for CPD. 1st quartile: top1 - mean effect size = 24.40%, 95% CI effect size = [18.20%, 30.61%], p = 1.124e-07; top2 - mean effect size = 33.12%, 95% CI effect size = [27.39%, 38.86%], p = 1.250e-10; top3 - mean effect size = 34.51%, 95% CI effect size = [29.51%, 39.51%], p = 3.980e-12; top4 - mean effect size = 34.70%, 95% CI effect size = [29.53%, 39.87%], p = 6.885e-12; top5 - mean effect size = 35.63%, 95% CI effect size = [30.26%, 41.01%], p = 8.878e-12. 2nd quartile: top1 - mean effect size = 32.96%, 95% CI effect size = [27.42%, 38.49%], p = 7.032e-11; top2 - mean effect size = 38.79%, 95% CI effect size = [33.73%, 43.84%], p = 4.837e-13; top3 - mean effect size = 40.45%, 95% CI effect size = [35.58%, 45.33%], p = 1.003e-13; top4 - mean effect size = 42.43%, 95% CI effect size = [37.62%, 47.24%], p = 2.827e-14; top5 - mean effect size = 42.46%, 95% CI effect size = [37.67%, 47.25%], p = 2.595e-14. 3rd quartile: top1 - mean effect size = 19.99%, 95% CI effect size = [14.47%, 25.51%], p = 3.175e-07; top2 - mean effect size = 22.78%, 95% CI effect size = [16.36%, 29.20%], p = 4.413e-07; top3 - mean effect size = 23.65%, 95% CI effect size = [16.71%, 30.58%], p = 8.273e-07; top4 - mean effect size = 23.67%, 95% CI effect size = [16.39%, 30.96%], p = 1.708e-06; top5 - mean effect size = 23.43%, 95% CI effect size = [16.17%, 30.70%], p = 1.918e-06. 4th quartile: top1 - mean effect size = - 5.96%, 95% CI effect size = [-9.80%, -2.12%], p = 5.834e-03; top2 - mean effect size = -6.32%, 95% CI effect size = [-10.24%, - 2.41%], p = 4.364e-03; top3 - mean effect size = -6.82%, 95% CI effect size = [-10.77%, -2.86%], p = 2.630e-03; top4 - mean effect size = -6.70%, 95% CI effect size = [-10.16%, -3.23%], p = 9.588e-04; top5 - mean effect size = -6.46%, 95% CI effect size = [-9.99%, -2.92%], p = 1.601e-03. **(b)** Accuracy by rank and quartile for StatAtlas. 1st quartile: top1 - mean effect size = 26.44%, 95% CI effect size = [23.56%, 29.32%], p = 1.265e-14; top2 - mean effect size = 27.31%, 95% CI effect size = [24.42%, 30.20%], p = 6.743e-15; top3 - mean effect size = 26.80%, 95% CI effect size = [23.62%, 29.98%], p = 7.078e-14; top4 - mean effect size = 27.89%, 95% CI effect size = [23.77%, 32.02%], p = 6.028e-12; top5 - mean effect size = 26.03%, 95% CI effect size = [21.95%, 30.11%], p = 1.840e-11. 2nd quartile: top1 - mean effect size = 27.71%, 95% CI effect size = [25.22%, 30.20%], p = 2.244e-16; top2 - mean effect size = 27.48%, 95% CI effect size = [24.62%, 30.35%], p = 5.028e-15; top3 - mean effect size = 26.09%, 95% CI effect size = [23.60%, 28.58%], p = 7.857e-16; top4 - mean effect size = 25.60%, 95% CI effect size = [23.16%, 28.04%], p = 7.552e-16; top5 - mean effect size = 24.67%, 95% CI effect size = [21.88%, 27.46%], p = 2.711e-14. 3rd quartile: top1 - mean effect size = 19.75%, 95% CI effect size = [16.14%, 23.36%], p = 2.053e-10; top2 - mean effect size = 20.62%, 95% CI effect size = [16.74%, 24.50%], p = 3.608e-10; top3 - mean effect size = 21.18%, 95% CI effect size = [17.93%, 24.43%], p = 6.459e-12; top4 - mean effect size = 20.90%, 95% CI effect size = [17.37%, 24.44%], p = 4.479e-11; top5 - mean effect size = 21.27%, 95% CI effect size = [17.20%, 25.34%], p = 4.985e-10. 4th quartile:top1 - mean effect size = 12.84%, 95% CI effect size = [11.00%, 14.68%], p = 1.557e-12; top2 - mean effect size = 10.57%, 95% CI effect size = [8.72%, 12.42%], p = 8.887e-11; top3 - mean effect size = 10.05%, 95% CI effect size = [8.53%, 11.57%], p = 4.861e-12; top4 - mean effect size = 9.44%, 95% CI effect size = [7.84%, 11.03%], p = 4.475e-11; top5 - mean effect size = 7.53%, 95% CI effect size = [5.83%, 9.23%], p = 1.037e-08. **(c)** Accuracy by rank and quartile for CRF ID. 1st quartile: top1 - mean effect size = 19.97%, 95% CI effect size = [15.62%, 24.32%], p = 8.294e-09; top2 - mean effect size = 16.07%, 95% CI effect size = [12.47%, 19.67%], p = 1.329e-08; top3 - mean effect size = 11.56%, 95% CI effect size = [7.71%, 15.40%], p = 6.574e-06; top4 - mean effect size = 8.63%, 95% CI effect size = [4.70%, 12.55%], p = 2.912e-04; top5 - mean effect size = 7.24%, 95% CI effect size = [3.38%, 11.09%], p = 1.342e-03. 2nd quartile: top1 - mean effect size = 21.42%, 95% CI effect size = [11.84%, 31.00%], p = 2.412e-04; top2 - mean effect size = 20.18%, 95% CI effect size = [9.41%, 30.94%], p = 1.360e-03; top3 - mean effect size = 15.61%, 95% CI effect size = [4.96%, 26.26%], p = 8.945e-03; top4 - mean effect size = 14.52%, 95% CI effect size = [4.12%, 24.92%], p = 1.220e-02; top5 - mean effect size = 13.62%, 95% CI effect size = [3.71%, 23.53%], p = 1.343e-02. 3rd quartile: top1 - mean effect size = 21.73%, 95% CI effect size = [15.50%, 27.96%], p = 5.751e-07; top2 - mean effect size = 16.31%, 95% CI effect size = [9.15%, 23.47%], p = 1.789e-04; top3 - mean effect size = 12.68%, 95% CI effect size = [5.67%, 19.68%], p = 1.737e-03; top4 - mean effect size = 10.92%, 95% CI effect size = [4.19%, 17.64%], p = 4.178e-03; top5 - mean effect size = 9.21%, 95% CI effect size = [2.76%, 15.66%], p = 1.027e- 02. 4th quartile: top1 - mean effect size = 10.63%, 95% CI effect size = [8.26%, 13.00%], p = 8.450e-09; top2 - mean effect size = 7.58%, 95% CI effect size = [5.54%, 9.62%], p = 2.031e-07; top3 - mean effect size = 6.54%, 95% CI effect size = [4.95%, 8.12%], p = 3.564e-08; top4 - mean effect size = 5.63%, 95% CI effect size = [4.15%, 7.10%], p = 1.420e-07; top5 - mean effect size = 4.87%, 95% CI effect size = [3.62%, 6.12%], p = 9.202e-08.

**Supplement 6:**
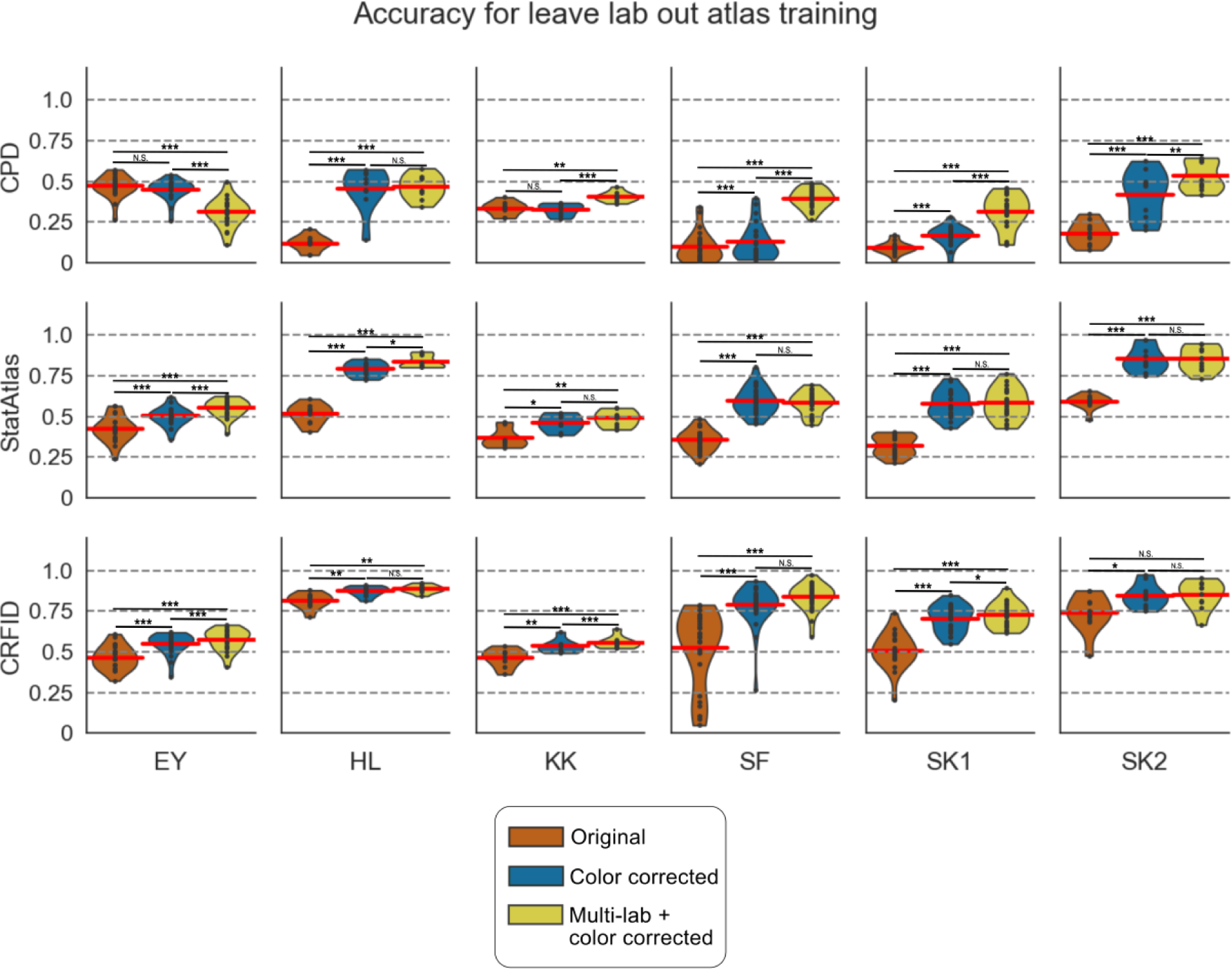
Top ranked assignment accuracy for leave-one-lab-out training. For each dataset, we train using the full corpus of data excluding all data from the dataset being tested. We then report the test accuracy for the dataset that was left out. All statistics are conducted with a paired t-test. Significance is reported using a Bonferroni correction with the convention of * for p<0.05, ** for p<0.01, and *** for p<0.001. **(a)** Accuracy by dataset for CPD. EY: original (10 worms) no color correction versus with leave-lab-out with color correction - mean effect size = -2.00%, 95% CI effect size = [-3.60%, -0.39%], p = 2.491e-02; original with no color correction versus multi-lab with color correction - mean effect size = -15.97%, 95% CI effect size = [-19.26%, -12.68%], p = 7.764e-09; leave-lab-out with color correction versus multi-lab with color correction - mean effect size = -13.97%, 95% CI effect size = [-17.10%, -10.84%], p = 3.041e-08. HL: original (10 worms) no color correction versus with leave-lab-out with color correction - mean effect size = 33.75%,95% CI effect size = [27.57%, 39.93%], p = 6.815e-06; original with no color correction versus multi-lab with color correction - mean effect size = 34.68%,95% CI effect size = [31.18%, 38.18%], p = 6.960e-08; leave- lab-out with color correction versus multi-lab with color correction - mean effect size = 0.93%,95% CI effect size = [-6.29%, 8.15%], p = 8.137e-01. KK: original (10 worms) no color correction versus with leave-lab-out with color correction - mean effect size = -0.67%,95% CI effect size = [-3.01%, 1.67%], p = 6.075e-01; original with no color correction versus multi-lab with color correction - mean effect size = 7.28%,95% CI effect size = [3.80%, 10.75%], p = 5.809e-03; leave-lab-out with color correction versus multi-lab with color correction - mean effect size = 7.95%,95% CI effect size = [5.26%, 10.63%], p = 8.722e-04. SF: original (10 worms) no color correction versus with leave-lab-out with color correction - mean effect size = 2.96%,95% CI effect size = [1.83%, 4.09%], p = 2.671e-05; original with no color correction versus multi-lab with color correction - mean effect size = 29.87%,95% CI effect size = [27.07%, 32.67%], p = 2.330e-17; leave-lab-out with color correction versus multi-lab with color correction - mean effect size = 26.91%,95% CI effect size = [23.62%, 30.20%], p = 1.165e-14. SK1: original (10 worms) no color correction versus with leave-lab-out with color correction - mean effect size = 7.39%,95% CI effect size = [5.08%, 9.70%], p = 5.516e-06; original with no color correction versus multi-lab with color correction - mean effect size = 22.22%,95% CI effect size = [17.35%, 27.08%], p = 3.335e-08; leave-lab-out with color correction versus multi-lab with color correction - mean effect size = 14.83%,95% CI effect size = [9.00%, 20.65%], p = 8.723e-05. SK2: original (10 worms) no color correction versus with leave-lab-out with color correction - mean effect size = 23.88%,95% CI effect size = [17.33%, 30.44%], p = 6.968e-05; original with no color correction versus multi-lab with color correction - mean effect size = 35.29%,95% CI effect size = [31.00%, 39.58%], p = 7.940e-08; leave-lab-out with color correction versus multi-lab with color correction - mean effect size = 11.41%,95% CI effect size = [5.67%, 17.15%], p = 4.398e-03. **(b)** Accuracy by dataset for StatAtlas. EY: original (10 worms) no color correction versus with leave-lab-out with color correction - mean effect size = 7.63%,95% CI effect size = [5.54%, 9.72%], p = 6.863e-07; original with no color correction versus multi-lab with color correction - mean effect size = 12.85%,95% CI effect size = [10.79%, 14.91%], p = 1.080e-10; leave-lab-out with color correction versus multi-lab with color correction - mean effect size = 5.22%,95% CI effect size = [4.03%, 6.41%], p = 4.261e-08. HL: original (10 worms) no color correction versus with leave-lab-out with color correction - mean effect size = 27.40%,95% CI effect size = [23.34%, 31.46%], p = 1.361e-06; original with no color correction versus multi-lab with color correction - mean effect size = 31.95%,95% CI effect size = [27.70%, 36.20%], p = 5.955e-07; leave-lab-out with color correction versus multi-lab with color correction - mean effect size = 4.55%,95% CI effect size = [1.93%, 7.17%], p = 1.123e-02. KK: original (10 worms) no color correction versus with leave-lab-out with color correction - mean effect size = 9.27%,95% CI effect size = [4.06%, 14.49%], p = 1.268e-02; original with no color correction versus multi-lab with color correction - mean effect size = 12.22%,95% CI effect size = [7.57%, 16.87%], p = 1.719e-03; leave-lab-out with color correction versus multi-lab with color correction - mean effect size = 2.94%,95% CI effect size = [0.74%, 5.15%], p = 4.095e-02. SF: original (10 worms) no color correction versus with leave-lab-out with color correction - mean effect size = 24.01%,95% CI effect size = [20.07%, 27.95%], p = 7.939e-12; original with no color correction versus multi-lab with color correction - mean effect size = 22.85%,95% CI effect size = [20.21%, 25.50%], p = 3.298e- 15; leave-lab-out with color correction versus multi-lab with color correction - mean effect size = -1.16%,95% CI effect size = [- 4.18%, 1.87%], p = 4.601e-01. SK1: original (10 worms) no color correction versus with leave-lab-out with color correction - mean effect size = 25.56%,95% CI effect size = [23.05%, 28.08%], p = 3.666e-14; original with no color correction versus multi-lab with color correction - mean effect size = 25.88%,95% CI effect size = [22.85%, 28.90%], p = 8.466e-13; leave-lab-out with color correction versus multi-lab with color correction - mean effect size = 0.31%,95% CI effect size = [-1.52%, 2.15%], p = 7.414e-01. SK2: original (10 worms) no color correction versus with leave-lab-out with color correction - mean effect size = 26.04%,95% CI effect size = [21.35%, 30.72%], p = 2.302e-06; original with no color correction versus multi-lab with color correction - mean effect size = 26.06%,95% CI effect size = [21.88%, 30.25%], p = 8.712e-07; leave-lab-out with color correction versus multi-lab with color correction - mean effect size = 0.03%,95% CI effect size = [-1.57%, 1.63%], p = 9.738e-01. **(c)** Accuracy by dataset for CRFID. EY: original (10 worms) no color correction versus with leave-lab-out with color correction - mean effect size = 8.51%,95% CI effect size = [5.94%, 11.08%], p = 2.641e-06; original with no color correction versus multi-lab with color correction - mean effect size = 10.97%,95% CI effect size = [8.32%, 13.62%], p = 9.944e-08; leave-lab-out with color correction versus multi-lab with color correction - mean effect size = 2.46%,95% CI effect size = [1.33%, 3.60%], p = 4.055e-04. HL: original (10 worms) no color correction versus with leave-lab-out with color correction - mean effect size = 5.86%,95% CI effect size = [2.81%, 8.91%], p = 6.772e-03; original with no color correction versus multi-lab with color correction - mean effect size = 7.28%,95% CI effect size = [4.05%, 10.50%], p = 2.776e-03; leave-lab-out with color correction versus multi-lab with color correction - mean effect size = 1.42%,95% CI effect size = [0.26%, 2.58%], p = 4.977e-02. KK: original (10 worms) no color correction versus with leave-lab-out with color correction - mean effect size = 7.07%,95% CI effect size = [4.27%, 9.87%], p = 2.133e-03; original with no color correction versus multi-lab with color correction - mean effect size = 9.42%,95% CI effect size = [6.72%, 12.12%], p = 3.264e-04; leave-lab-out with color correction versus multi-lab with color correction - mean effect size = 2.35%,95% CI effect size = [1.76%, 2.93%], p = 1.356e-04. SF: original (10 worms) no color correction versus with leave-lab-out with color correction - mean effect size = 26.05%,95% CI effect size = [18.80%, 33.29%], p = 2.164e-07; original with no color correction versus multi-lab with color correction - mean effect size = 31.28%,95% CI effect size = [23.27%, 39.28%], p = 5.119e- 08; leave-lab-out with color correction versus multi-lab with color correction - mean effect size = 5.23%,95% CI effect size = [1.22%, 9.24%], p = 1.705e-02. SK1: original (10 worms) no color correction versus with leave-lab-out with color correction - mean effect size = 19.66%,95% CI effect size = [15.19%, 24.13%], p = 5.897e-08; original with no color correction versus multi-lab with color correction - mean effect size = 21.75%,95% CI effect size = [17.54%, 25.97%], p = 4.768e-09; leave-lab-out with color correction versus multi-lab with color correction - mean effect size = 2.09%,95% CI effect size = [0.66%, 3.52%], p = 1.015e-02. SK2: original (10 worms) no color correction versus with leave-lab-out with color correction - mean effect size = 10.61%,95% CI effect size = [3.88%, 17.34%], p = 1.521e-02; original with no color correction versus multi-lab with color correction - mean effect size = 10.91%,95% CI effect size = [3.73%, 18.08%], p = 1.803e-02; leave-lab-out with color correction versus multi-lab with color correction - mean effect size = 0.30%,95% CI effect size = [-3.51%, 4.11%], p = 8.847e-01.

**Supplement 7:**
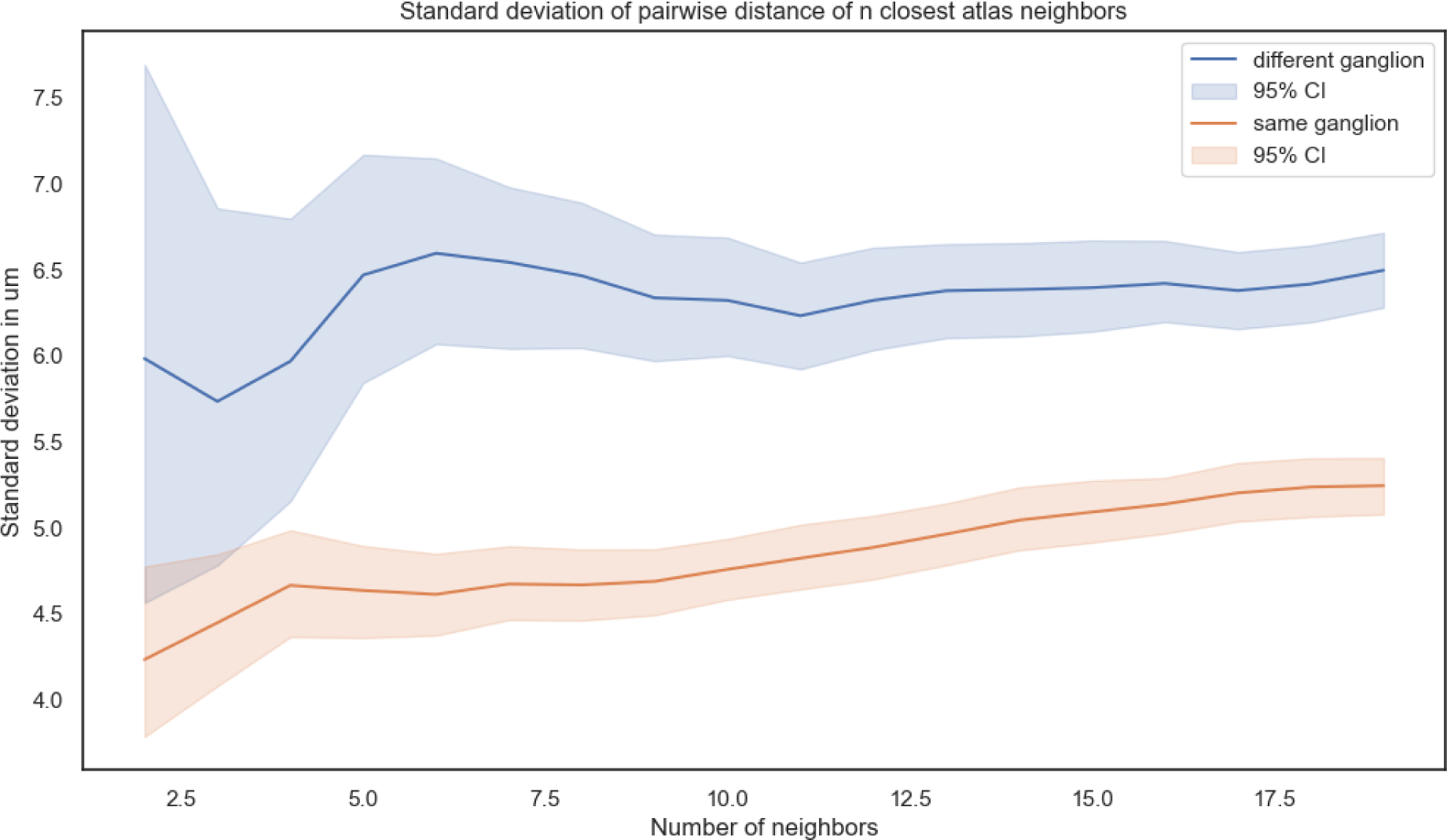
Standard deviation of pairwise distance of *n* closest atlas neighbors.

